# Effects of Force Fields on the Conformational and Dynamic Properties of Amyloid β(1-40) Dimer Explored by Replica Exchange Molecular Dynamics Simulations

**DOI:** 10.1101/210286

**Authors:** Charles R. Watts, Andrew Gregory, Cole Frisbie, Sándor Lovas

**Affiliations:** Department of Neurosurgery, Mayo Clinic, College of Medicine, Rochester, MN, 55905, USA.; Department of Neurosurgery, Mayo Clinic Health System, La Crosse, WI, 54601, USA.; Department of Biomedical Sciences, Creighton University, Omaha, NE, 61718, USA.

**Keywords:** Amyloid β, Dimerization, REMD simulations, Force Fields, Loosely packed anti-parallel β-sheet dimer, Inherently Disordered Peptide, Alzheimer’s Disease, Cerebral Amyloid Angiopathy, Statistical Analysis

## Abstract

Alzheimer’s disease is histologically marked by fibrils of Amyloid beta (Aβ) peptide within the extracellular matrix. Fibrils themselves are benign compared to the cytotoxicity of the oligomers and pre-fibrillary aggregates. The conformational space and structural ensembles of Aβ peptides and their oligomers in solution are inherently disordered and proven to be challenging to study. Optimum force field selection for molecular dynamics (MD) simulations and the biophysical relevance of results are still unknown. We compared the conformational space of the Aβ(1–40) dimers by 300 ns replica exchange MD simulations at physiological temperature (310 K) using: the AMBER-ff99sb-ILDN, AMBER-ff99sb*-ILDN, AMBER-ff99sb-NMR, and CHARMM22* force fields. Statistical comparisons of simulation results to experimental data and previously published simulations utilizing the CHARMM22* and CHARMM36 force fields were performed. All force fields yield sampled ensembles of conformations with collision cross sectional areas for the dimer that are statistically significantly larger than experimental results. All force fields, with the exception of AMBER-ff99sb-ILDN (8.8±6.4%) and CHARMM36 (2.7±4.2%), tend to overestimate the α-helical content compared to experimental CD (5.3±5.2%). Using the AMBER-ff99sb-NMR force field resulted in the greatest degree of variance (41.3±12.9%). Except for the AMBER-ff99sb-NMR force field, the others tended to under estimate the expected amount of β-sheet and over estimate the amount of turn/bend/random coil conformations. All force fields, with the exception AMBER-ff99sb-NMR, reproduce a theoretically expected β-sheet-turn-β-sheet conformational motif, however, only the CHARMM22* and CHARMM36 force fields yield results compatible with collapse of the central and C-terminal hydrophobic cores from residues 17-21 and 30-36. Although analyses of essential subspace sampling showed only minor variations between force fields, secondary structures of lowest energy conformers are different.

## Introduction

Amyloid β(Aβ) peptides are the product of the proteolytic cleavage of amyloid precursor protein by β- and γ- secretases. With the exception of several genetic mutations, glycosylation, and minor isoforms, the two dominant isoforms found in cerebrospinal fluid and cerebral cortical tissue are Aβ(1-40) and Aβ(1-42). These two isoforms play an important role in the Alzheimer’s Dementia (AD), and Cerebral Amyloid Angiopathy (CAA). The amyloid hypothesis of disease causation postulates that the development of these diseases is related to an imbalance between production and clearance of Aβ peptides that results in oligomerization and eventual Aβ fibril formation that is the histological hallmark of both AD and CAA.^1-10^ Despite the histological and diagnostic importance of Aβ fibrils, the aggregates and pre-fibril forms of Aβ peptides that are a result of Aβ oligomerization have been shown to be more cytotoxic that the fibrils and may play a crucial role in disease development.^11,12^

Among the major forms of Aβ, Aβ(1-40) is more soluble than Aβ(1-42) and has much slower kinetics of oligomer and fibril formation than Aβ(1-42).^13^ Aβ(1-40) fibrils are more common in the walls of cerebral vessels while Aβ(1-42) fibrils are more common in cerebral cortical tissue. The predominance of Aβ(1-42) fibrils in cerebral cortical tissue does not necessarily correlate with causation of AD or CAA given the finding that Aβ(1-40) is ten times more abundant in cerebrospinal fluid than Aβ(1-42) ^14-16^

The shared primary structural feature of Aβ peptides is a central hydrophobic core (CHC) from residues Leu^17^ to Ala^21^ and a C-terminal hydrophobic core from Ala^30^ to Val^36^ with an N-terminal hydrophilic region from Asp^1^ to Lys^16^ and central hydrophilic region from Glu^22^ to Gly^29^. X-ray diffraction, solid state NMR, other spectroscopic methods and molecular dynamics (MD) simulations have demonstrated that collapse of CHC and residues 30-36, leads to the formation of a β-strand-loop-β-strand motif that is important for oligomerization and fibril formation^17-20^ This β-strand-loop-β-strand motif is common in other amyloid forming peptides and appears to arise from the propensity of peptide chains to form H-bonds between backbone atoms with packing for the β-sheets into higher order structures is a consequence of specific side chain interactions.^21^ Although the last four and six C-terminal residues of Aβ(1-40) and Aβ(1-42), respectively, undoubtedly play a role in isoform solution and biologic behavior, current MD simulation studies have indicated that their conformation are largely disordered and they do not play a role in the β-strand-loop-β-strand motif formation.^22-26^

The presence of the β-strand-loop-β-strand motif appears to be relatively constant across most amyloid forming peptides in an aggregated state, but their behavior in solution is inherently disordered and their structural determination is more difficult.^19^,^20,27,28^ There are also significant differences between the solution conformations sampled by Aβ(1-40) monomer and that of its higher order oligomers. The NMR solution structure of Aβ(1-40) in 50 mM NaCl aqueous solution at neutral pH and 288 K (PDB ID: 2LFM) demonstrates that the N- and C-terminal regions are largely unstructured or transitionally structured with a 3_10_-helix from residues His^13^ to Asp^23^ spanning the CHC from residues 17 to 21.^29^ This behavior may also hold for other forms of Aβ monomers given a recently published MD simulation of Aβ(10-40) using the CHARMM22/cmap and CHARMM36 force fields demonstrated a similar helical motif with excellent agreement with the previously published ^3^ J_HNH_-coupling and RDC constants from the 2LFM solution structure.^30^

The lowest order Aβ oligomer that appears to be biologically active is the dimer having been identified in the cerebral tissue of AD patients.^16^ Covalently linked synthetic Aβ dimers have also been used to demonstrate rapid oligomerization and fibril formation as well as the development of hyper-phosphorylated tau protein in the cerebral cortex tissue of animal models.^31-33^ Aβ dimer, therefore, becomes the simplest and most computationally accessible oligomer to study. Two recently published replica exchange dynamics (REMD) simulations studied the conformations of Aβ(1-40) dimers.^23,26^ The study by Tarus and colleagues^23^ utilized the CHARMM22* force field and analyzed the data at 315 K, NVT ensemble. Their model has four β-strand regions from residues 3 to 6, 10 to 12, 17 to 21, and 31 to 36 separated by loop/turn regions from residues 7 to 9, 13 to 15, and 23 to 29. There were transiently populated (5 to 17%) helical regions occurring within the N-terminus, CHC, and residues 22 to 29 and 30 to 38. The tertiary structure of the monomers and quaternary structure of the dimer were largely disordered. Our recently published model of Aβ(1-40)^26^, using the CHARMM36 force field and the trajectory analyzed at 310 K, NPT ensemble, has β-sheets from residues 17 to 21 and 30 to 33 separated by a mixed β-turn/loop conformation from residues 22 to 29. This conformation corresponds to formation of the β-strand-loop-β-strand motif caused by hydrogen bonding between the two chains and loose collapse and packing of the hydrophobic side chains of the central hydrophobic and C-terminal hydrophobic cores. Similar to the results of Tarus and colleagues the resulting tertiary and quaternary structures were largely disordered.^23,26^

Although it would appear that the CHARMM36 results are better at reproducing the expected β-strand-loop-β-strand motif, recent experiments utilizing the CHARMM36 force field on a series of inherently disordered peptides has demonstrated problems with increased sampling of the α_L_-helical region of ϕ/ψ dihedral angle space resulting in generation of unrealistic conformations.^34^ This phenomenon of increased α_L_-helical sampling was not observed in our previous study and it is unclear if the highly polar nature of RS, (AAQAA)_3_, and HEWL19 peptides played a significant role in the findings of Rauscher *et al*.^34^ Despite multiple MD simulations on Aβ peptides, it is neither clear what force field should be used for simulations nor how to compare and interpret the results of simulations completed utilizing different force fields.^30,35-39^

Herein, we utilize statistically robust methodologies comparing the conformational and dynamic properties of Aβ(1-40) dimer as a function of force field selection using REMD simulations in TIP3P water with the AMBER-ff99sb-ILDN, AMBER-ff99sb*-ILDN, AMBER-ff99sb-NMR, and CHARMM22* force fields.^40-48^ Comparisons are made to existing circular dichroism spectropolarimetry (CD), IM-MS collision cross section measurements (CCS), and the previously published dimer simulation of Tarus and colleagues^23^ using the CHARMM22* force field and our previous results with CHARMM36 force fields.^26^

## Methods

## REMD Simulations

The dimer structure was created by randomly associating two of the first NMR structure of Aβ(1-40) (PDB ID.: 2LFM) such that their center of mass distance was 1 nm as described in detail in our previous work.^26^ For the REMD simulations four separate force fields as implemented in the GROMACS version 4.6.4 were used: AMBER-ff99sb-ILDN (AMBER-ILDN), AMBER-ff99sb*-ILDN (AMBER-ILDN*), AMBER-ff99sb-NMR (AMBER-NMR), and CHARMM22*.^40-48^ The initial dimer structure was solvated in a truncated octahedron of 203.02 nm^3^ with 6235 TIP3P water molecules.^49^ The system was neutralized with 24 Na^+^ and 18 Cl^−^ ions and the final concentration of the NaCl was set to 150 mM. The system was subjected to 1000 steps steepest descent energy minimization and then to 100 ps NVT simulation at 300 K so that the position of the dimer was constrained at the center of the octahedron with a force constant of 1000 kJ•mol^−1^. The unrestrained system was then submitted to a 500 ps NPT simulation at 1 bar pressure and 300 K temperature to allow for adequate system equilibration and relaxation. The integration step was 2 fs, the LINCS^50^ algorithm was used to constrain all bonds to their correct length with a warning angle of 30°; The peptide and the solvent with ions were coupled to separate temperature baths with a relaxation constant of 0.1 ps; The peptide and the solvent with ions were coupled separately to constant pressure using Berendsen scaling with a relaxation constant of 1.0 ps and 4.5 × 10^−5^ bar^−1^ isothermal compressibility. ^51^ The temperature was controlled by the stochastic velocity-rescaling method of Bussi *et al*.^52^ The long-range electrostatic interactions were calculated using the PME method with 1.2 nm and 1.0 nm cutoff distance and the Fourier spacing was 0.15 nm. For the calculations of van der Waals interactions the short-range and long-range cutoffs, respectively, were 1.0 and 1.2 nm for the CHARMM22* force field and 0.7 and 0.9 nm for the AMBER force fields. Dispersion correction was applied to both the energy and the pressure. The 300 ns NPT REMD simulations were performed with 70 replicas of the initial structure for a total replica simulation time of 21,000 ns for each force field and a total simulation time of 84,000 ns utilizing all four force fields. The exchange temperatures were obtained using the temperature predictor web server (http://folding.bmc.uu.se/remd/) with the lower and upper limits of 290 K and 470 K, respectively, and exchange probability of 0.2, occurring every 2 ps, only exchanging replicas with neighboring temperatures.^53-55^ The dimer and solvent with ions were separately coupled to a 1 bar Parrinello-Rahman barostat and the temperature was maintained by the stochastic velocity-rescaling method.^52,56,57^ The non-bonded parameters were as above.

The GROMACS program writes trajectory files from REMD simulations that are continuous with respect to ensemble but not with respect to simulations time. In order to be able to follow the replicas of interest through constant temperature space, the REMD trajectories were demultiplexed for constant temperature and continuous time using the g_*trjcat* module and the demultiplexing matrix which was generated by the *demux.pl perl* script (www.gromacs.org/Documentation).

## Trajectory Analysis

### Determination of convergence

A sampling frequency of 0.1 ns was used for the analysis. To determine if the dimer system had achieved a thermodynamic equilibrium, an essential dynamics analysis of the trajectory was performed.^58,59^ The covariance matrix for the Cα-atoms was calculated using the *g_covar* module of GROMACS, the eigenvectors were used to calculate the backbone configurational entropy as a function of system temperature for all of the replicas from 290 K to 470 K.^60^ The statistical significance of convergence of the trajectories was obtained by calculation of the root mean square inner product (RMSIP) over the eigenvectors of the Cα-atoms with the exception of the very first *N*- and last *C*-terminal residues.^58,61,62^

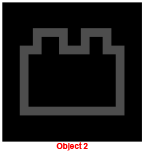

The eigenvectors η_i_ and η_j_ represent subparts of the trajectory of increasing time length taken from the beginning and the end of the simulation to prevent dynamic autocorrelation; n is the number of eigenvectors. RMSIP values range from 0 to 1; the value is 1 if the sampled subspaces are identical and 0 if they are orthogonal. Assuming that the computed covariance matrix is a reasonable estimate of one calculated at infinite time, the subpart eigenvectors should converge in an asymptotic fashion with long enough calculations.^62^ Values of RMSIP ≥ 0.6 are considered good convergence while RMSIPs ≥ 0.8 are considered excellent.^58^ Due to the increased sampling of α_L_-helical ϕ/ψ dihedral angle space noted by Rauscher *et al*.^34^, the ϕ-ψ dihedral angles of residues (2-39) of both chains were examined as a function of time by dividing the trajectory into 6 different intervals: 0 to 50 ns, 50 to 100 ns, 100 to 150 ns, 150 to 200 ns, 200 to 250 ns, and 250 to 300 ns. The sampled ϕ-ψ dihedral angles were extracted using the *g_rama* utility of GRO-MACS and plotted as a surface using the *plot3D* module of *R version 3.3.1*.^63,64^ Attention was given to population changes in the respective regions of ϕ-ψ.

### Biophysical Properties

The second nematic order parameter (*P*_2_), radius of gyration (R_g_), collision cross sectional area (CCS), fraction of populated secondary structure, and Cα-trace root mean square fluctuations (RMSF) were calculated, with the *Wordom* program, the *g_gyration* utility of GROMACS, *mobcal, do_dssp,* and *g_rmsf utilities* of GROMACS, respectively.^65-68^ A detailed description of the implementation of these methods can be found in our previous work.^26^ ***Interactions.*** The solvent exposed surface area (SASA) of the dimer and each chain where calculated with *g_sasa* utility of GROMACS using the atomic radii of Lee and Richards, a probe radius of 0.14 nm, and 1000 points per sphere resolution.^69^ The interaction surface area (ISA) between the two chains to form the dimer was calculated as:

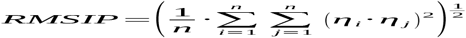

ISA was measured on a per residue basis (rISA) and normalized to values of the maximal solvent accessible surface area as determined from MD simulations on Ace-Ala-X-Ala-NH2.^26,70,71^ The intra- and inter- chain side chain-side chain and backbone-backbone contacts, H-bonds, and salt bridges were calculated with the *g_mdmat, g_hbond,* and *g_saltbr* utilities of GROMACS, respectively. Side chain-side chain and backbone-backbone contacts, were considered to occur if the inter-atomic distances between the heavy atoms of a residue were ≤0.5 nm. Intra- and inter-chain correlated motions were evaluated by calculating dynamic cross correlation matrices (DCCM) from the principal components of the Cα-trace covariance matrix using the method of Amadei *et al.* as implemented in the *Bio3d* module in *R version 3.3.1.*^64,72,73^ The results are displayed as a color coded matrix of Pearson correlation coeficients with a value of −1 indicated completely anticorrelated motions and a value of +1 indicating completely correlated motions.^74,75^ H-bonds were considered to occur if the donor acceptor radius was ≤0.35 nm and the donor-hydrogen-acceptor angle was ≤30°. The binding energy (ΔE_binding_) between the two monomers was calculated with the *g_mmpbsa* program as we described previously.^76-80,26^ Since the g_mmpbsa program does not calculate the entropy change upon binding, we only report ∆E_binding_.

### Conformational and Essential Subspace Analysis

The sampled conformations of the dimer and individual chain components were analyzed using the dihedral principal component analysis (dPCA) method.^81,82^ A detailed description of our method can be found in reference 26. The lowest energy conformations were identified by projecting the trajectories of the first three principal components (dPC1, dPC2, and dPC3) onto a three-dimensional free energy (∆G) landscape

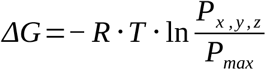

R is the universal gas constant, T is the temperature, x, y, and z, are calculated structural properties from the trajectory. Low energy regions were identified using grid based density clustering and representative conformations from each region extracted for further analysis.^26,83,84^ Grid based density methodologies have been shown to be robust for detecting clusters with non-spherical shape and high levels of noise in the data.^83-88^

To determine the overlap of the sampled essential subspace of different trajectories as a function of force field, the root mean square inner product was calculated;

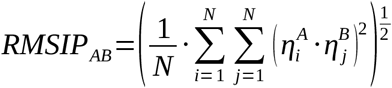

RMSIP_AB_ is the root mean square inner product comparing trajectories A and B, 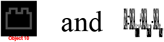 are the respective eigenvectors of the sampled essential subspace for the trajectories, and N is the total number of eigenvectors to be considered. ^58,61,62^ The RMSIP can then be normalized;

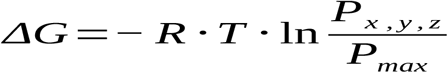

RMSIP_A1•A2_ and RMSIP_B1•B2_ are comparisons between the first and second sampled halves of trajectories A and B, respectively. Comparison the first and second halves of the trajectories corrects errors from autocorrelation and sampling. A value approaching unity indicates that the similarity between trajectories A and B is similar to that between the two halves of trajectories A and B. Lower values however, indicate differences between trajectories A and B that cannot be explained by sampling alone.^89^

### Statistical Analysis

A Detailed description of the statistical methods^90-101^ is in the Supporting Information (SI). The biophysical properties: *P*_2_, R_g_, CCS, fractions of populated secondary structure (β-sheet, α-helix, and turn/coil/bend), SASAs of the individual chains and dimer, ISAs, hydrophobic surface areas of the individual chains and dimer, hydrophilic surface areas of the individual chains and dimer, the change in hydrophobic, hydrophilic surface areas with dimerization, and the number of intra- and inter-chain backbone-backbone (BB-BB) and sidechain-sidechain (SC-SC) contacts were characterized for each force field with: mean, standard deviation, and 95% confidence interval (CI_95_). The homogeneity of variances were determined with Levene’s test and comparisons made with Welch’s t-test and Welch’s ANOVA where appropriate and *post-hoc* analysis with the Games-Howell test.^95-101^ In order to make comparisons with previously published data sets of limited size and ensure appropriate balance to Welch’s t-test and Welch’s ANOVA one of three methods was utilized. First, if only the maximum and minimum values were published, we made the assumption that the reported values represented the CI_95_ of the data and the midpoint represented the mean. Using these three values (min, max, and mean), the standard deviation was determined using a t-distribution with 2 degrees of freedom. The mean and t-distribution corrected standard deviation were used to bootstrap a normal distribution of 2000 values for analysis using the R program.^102^ Second, if several published values were available we again made the assumption that the reported maximum and minimum values represented the CI_95_ and the mean was calculated using all available data points. The standard deviation was determined using a t-distribution with n-1 degrees of freedom where n was the number of reported values. A normal distribution of 2000 values was then bootstrapped as discussed above. Third, if a mean and standard deviation were published and the respective values were obtained from a large database of samples (n>500), a normal distribution of 2000 values was bootstrapped without the t-distribution correction.

## Results

### Simulation Convergence

Because of the know difficulties in determining simulation convergence to a thermodynamics equilibrium, we used a multi-mode approach. The configurational entropy as a function of simulation time for the Cα-trace of the dimer systems at 300 K, 310 K, 314 K and 470 K are plotted in Figure 1. The configurational entropy of the AMBER-ILDN, AMBER-NMR, and CHARMM22* trajectories rise rapidly for all temperatures over the first 50 ns of the simulations while that of AMBER-ILDN* requires at least 75 ns of simulation time to reach a maximum for the 310 K trajectory and much longer periods of time for lower temperature trajectories as shown for 300 K. After the rapid rise in configurational entropy, all trajectories required at least 25 to 50 ns of simulation time in order to reach a stable plateau. The configurational entropy plateau value for the low temperatures (300 to 314 K) was similar for the AMBER-ILDN, and AMBER-NMR trajectories. Due to the long and gradual rise of the AMBER-ILDN* trajectory at 300 K, it is unclear if a similar plateau value would have been reached. There was also a significant decrease in Cα-trace configurational entropy comparing the 300 K trajectory to the 310 K, 314 K and 470 K trajectory plateaus for the CHARMM22* simulations.

**Figure 1.**
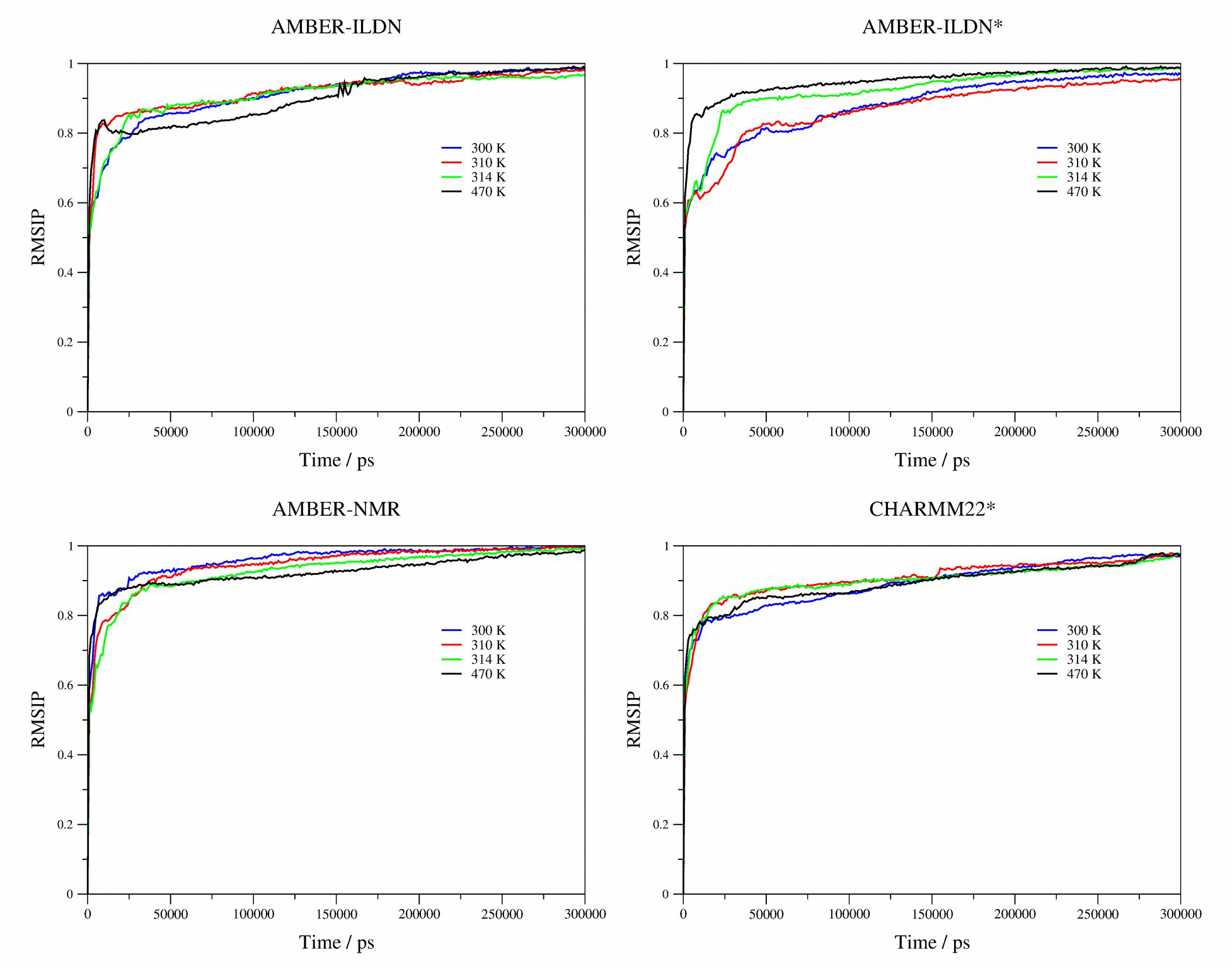
Configurational Cα-trace entropy of Aβ(1-40) dimer at 300 K, 310 K (physiological temperature), 314 K, and 470 K for AMBER-ILDN, AMBER-ILDN*, AMBER-NMR, and CHARMM22* force fields, demonstrating the 100 ns time necessary for system equilibration.

The statistical significance of variation in configurational space sampled by the trajectories as a function of time was determined by calculating the RMSIP for the eigenvectors of the Cα-trace. The results for 300 K, 310 K, 314 K, and 470 K are shown in Figure 2. The trajectories with the exception of AMBER-ILDN*, rise to a RMSIP > 0.8 within 50 ns. The RMSIP value AMBER-ILDN* does reach the >0.8 value indicative of a high level of convergence at a similar rate to the change in entropy as a function of time, requiring approximately 75 ns of simulation time. The RMSIP remains at a similar level of convergence from 100 to 300 ns indicating stable sampling of configurational space with a statistically stable degree of variation in the eigenvectors. We also compared RMSIPs of the trajectories of the individual chains to themselves and to each other (Figures S1 and S2 in the SI). RMSIPs for the individual chains reached asymptotic values of RMSIP>0.9 within 100 ns. The RMSIP between chain A and chain B converged to ~0.8 within similar time. The value of the RMSIP between chain A to chain B is only slightly lower than that of the individual chains and dimer demonstrating that there is significant shared conformational and dynamic similarity and the relationship between the two chains is not orthogonal. The similarity may be due to collapse of the hydrophobic core with more stable regions of β-sheet motif while the variability occurs within the flexible intervening loops and coils.

**Figure 2.**
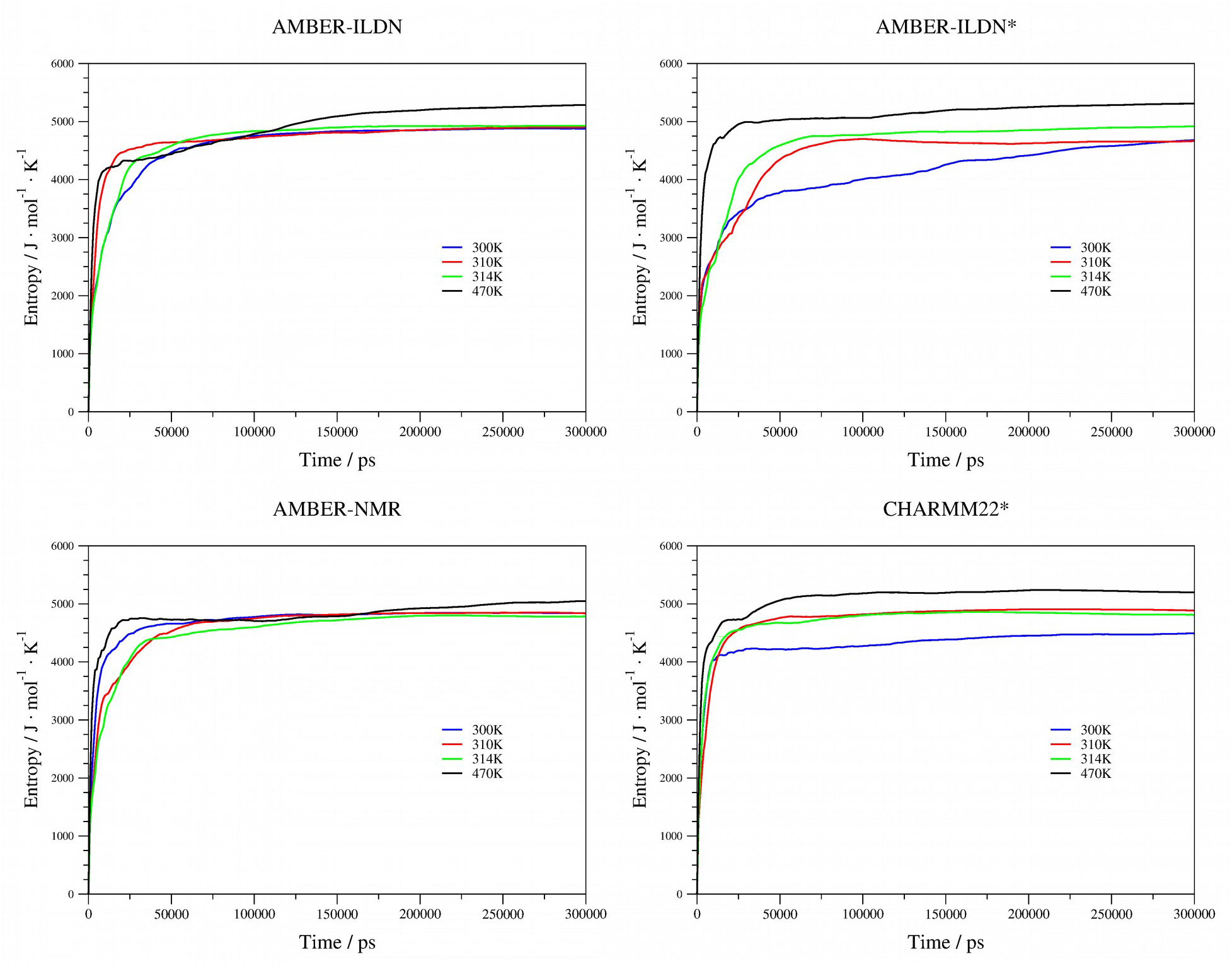
The Cα-trace root mean square inner product (RMSIP) for the eigenvectors of Aβ(1-40) dimer at 300 K, 310 K (physiological temperature), 314 K, and 470 K as a function of simulation time for AMBER-ILDN, AMBER-ILDN*, AMBER-NMR, and CHARMM22* force fields.

The dimer system at 310 K maintains a relatively stable secondary structure for the first 10 ns of the sampled trajectories for AMBER-ILDN, and CHARMM22* (Figure S3 in the SI). The most persistent secondary structural feature is a region of α-helix from residues 27 through 30 of Chain A and residues 18 through 20 of Chain B that gradually changes to bend, turn, and β-sheet configurations over the first 100 ns of the simulations. The AMBER-ILDN* trajectory shows persistent stability of the α-helical motif in the same residues over the first 25 ns of the simulation and continued high levels of α-helical sampling throughout the simulation. The AMBER-NMR trajectory is similar to AMBER-ILDN* for the first 25 ns however, after that time period, high levels of α-helical conformations are sampled across most residues through the remainder of the trajectory.

The sampled ϕ-ψ dihedrals at 310 K over 50 ns time frames demonstrate stable conformational space sampling from 100 to 300 ns. The AMBER-ILDN ϕ-ψ dihedrals demonstrate sampling of the β-sheet and α_R_-helical regions of conformational space (Figure 3). AMBER-ILDN*, AMBER-NMR, and CHARMM22* have an increased probability of sampled αR-helical conformational space with the AMBER-NMR sampling being heavily favored towards this region with a significant decrease in the β-sheet regions. AMBER-ILDN, AMBER-ILDN* and CHARMM22* all sample a small population of the α_L_-helical regions of conformational space while AMBER-NMR does not. In order to confirm that extended regions of α_L_-helical secondary structure were absent from the sampled conformational space, a detailed analysis of the *i-1* and *i+1* residues was done and demonstrated that if the ϕ-ψ dihedrals of residue *i* were within the α_L_-helical region of conformational space, the probability that either the *i-1* or *i+1* residue was within the same region of conformational space was < 0.01. Those residues whose ϕ-ψ dihedrals sampled α_L_-helical conformational space were, therefore, classified by the DSSP method as short three or four residue turns without hydrogen bonding.^68^ On the basis of configurational entropy, RMSIP, DSSP-calculated secondary structure as a function of time, and ϕ-ψ dihedral angles as a function of time, convergence of the simulations was likely reached prior to 100 ns and subsequent biophysical propert and conformational analysis was therefore performed using the 100 ns to 300 ns sampling period.

**Figure 3.**
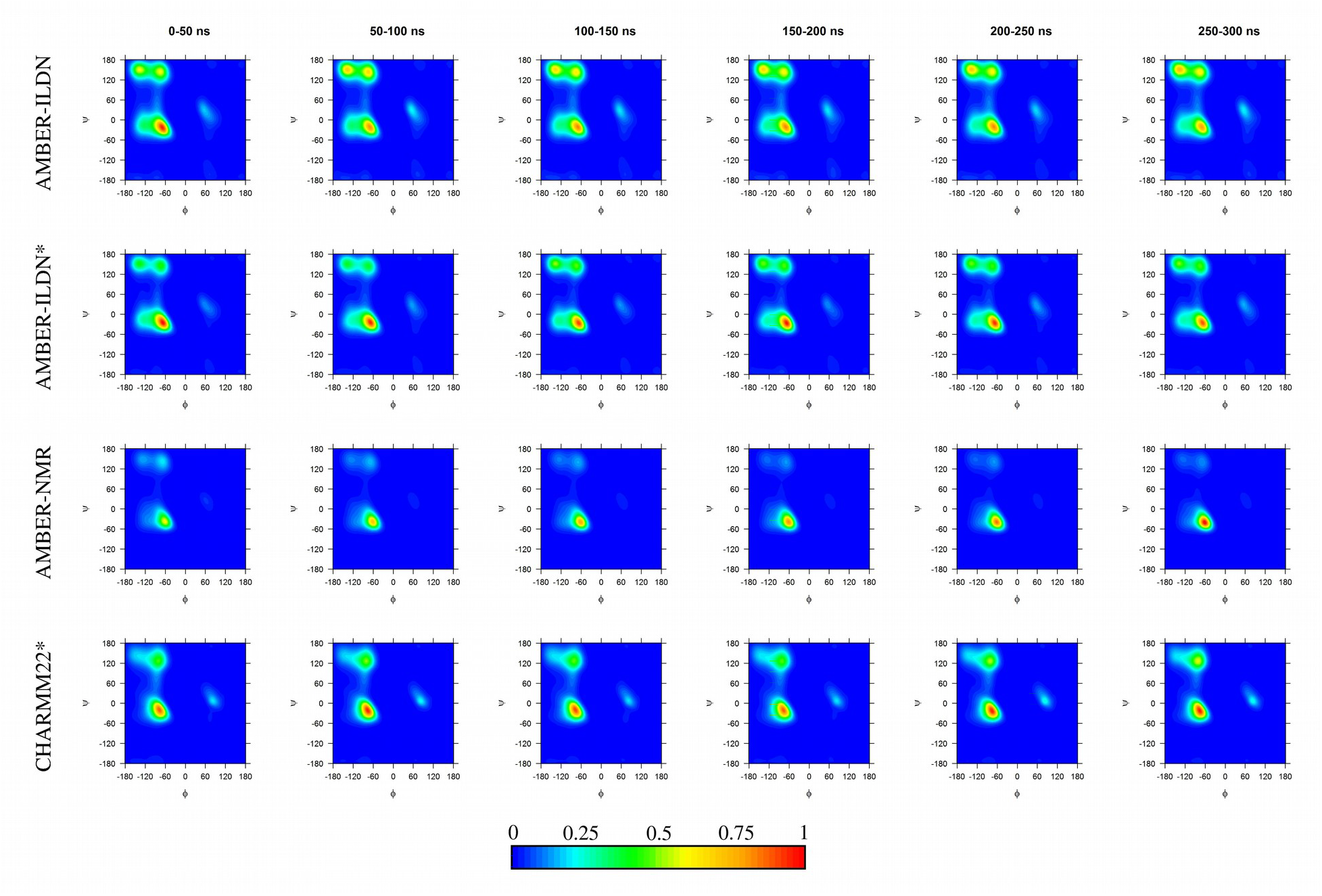
The sampled ϕ-ψ dihedral angles for residues 2 through 39 of Chain A and Chain B plotted at 50 ns intervals for AMBER-ILDN, AMBER-ILDN*, AMBER-NMR, and CHARMM22* force fields. The color scale bars represent the normalized global probability of a given ϕ-ψ dihedral angle being sampled during the respective time period.

### Biophysical Properties

Statistical comparisons were made between the calculated values of biophysical properties for each trajectory at 310 K, experimental values when available, and previously published dimer simulations as outlined in the supporting information (SI).^23,26^ The bootstrapped means and standard deviations for CCS and CD are given in Tables S1 and S2 in the SI.^105,106,104,110^ The results are collated in Table 1, box plots with whiskers representing the mean, standard deviation, and CI_95_ are shown in Figures S4 - S15 in the SI, and the results of Levene’s test and the associated F-statistics and p-values of the ANOVA are given in Table S3 of the SI.

**Table 1.**
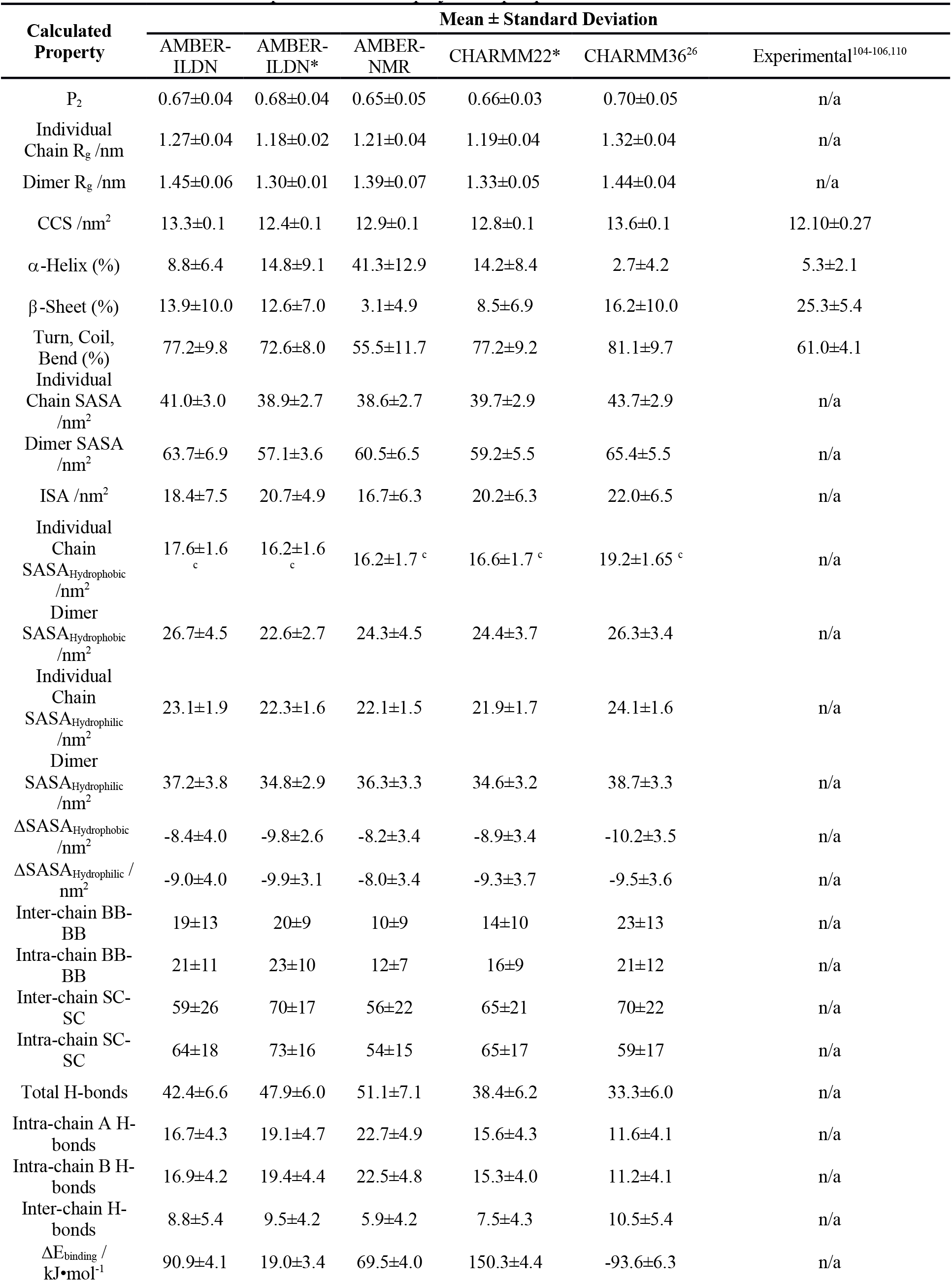
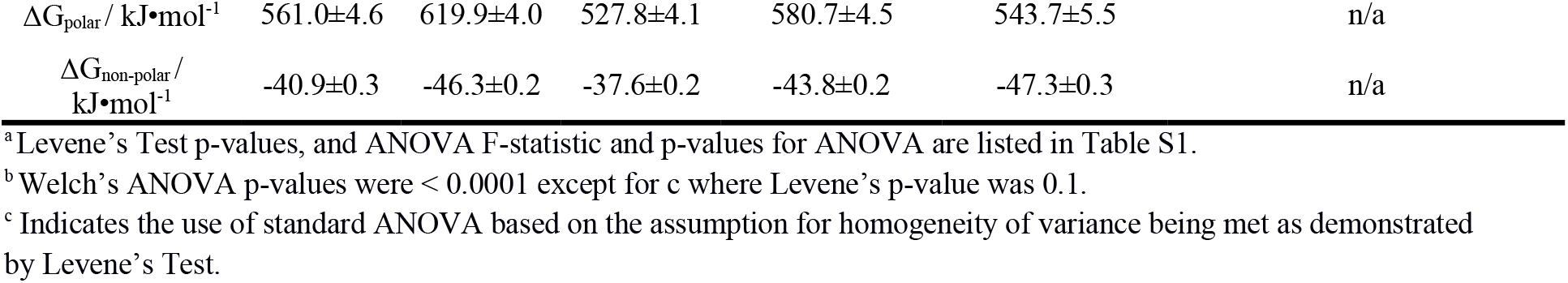
Calculated and experimental biophysical properties of dimers.^a,b^

| Calculated Property | Mean ± Standard Deviation | | | | | Experimental<sup>104-106,110</sup> |
| --- | --- | --- | --- | --- | --- | --- |
|  | AMBER-ILDN | AMBER-ILDN* | AMBER-NMR | CHARMM22* | CHARMM36<sup>26</sup> |  |
| P<sub>2</sub> | 0.67±0.04 | 0.68±0.04 | 0.65±0.05 | 0.66±0.03 | 0.70±0.05 | n/a |
| Individual Chain R<sub>g</sub> /nm | 1.27±0.04 | 1.18±0.02 | 1.21±0.04 | 1.19±0.04 | 1.32±0.04 | n/a |
| Dimer R<sub>g</sub> /nm | 1.45±0.06 | 1.30±0.01 | 1.39±0.07 | 1.33±0.05 | 1.44±0.04 | n/a |
| CCS /nm<sup>2</sup> | 13.3±0.1 | 12.4±0.1 | 12.9±0.1 | 12.8±0.1 | 13.6±0.1 | 12.10±0.27 |
| α-Helix (%) | 8.8±6.4 | 14.8±9.1 | 41.3±12.9 | 14.2±8.4 | 2.7±4.2 | 5.3±2.1 |
| β-Sheet (%) | 13.9±10.0 | 12.6±7.0 | 3.1±4.9 | 8.5±6.9 | 16.2±10.0 | 25.3±5.4 |
| Turn, Coil, Bend (%) | 77.2±9.8 | 72.6±8.0 | 55.5±11.7 | 77.2±9.2 | 81.1±9.7 | 61.0±4.1 |
| Individual Chain SASA /nm<sup>2</sup> | 41.0±3.0 | 38.9±2.7 | 38.6±2.7 | 39.7±2.9 | 43.7±2.9 | n/a |
| Dimer SASA /nm<sup>2</sup> | 63.7±6.9 | 57.1±3.6 | 60.5±6.5 | 59.2±5.5 | 65.4±5.5 | n/a |
| ISA /nm<sup>2</sup> | 18.4±7.5 | 20.7±4.9 | 16.7±6.3 | 20.2±6.3 | 22.0±6.5 | n/a |
| Individual Chain SASA<sub>Hydrophobic</sub> /nm<sup>2</sup> | 17.6±1.6<sub>c</sub> | 16.2±1.6<sub>c</sub> | 16.2±1.7<sup>c</sup> | 16.6±1.7<sup>c</sup> | 19.2±1.65<sup>c</sup> | n/a |
| Dimer SASA<sub>Hydrophobic</sub> /nm<sup>2</sup> | 26.7±4.5 | 22.6±2.7 | 24.3±4.5 | 24.4±3.7 | 26.3±3.4 | n/a |
| Individual Chain SASA<sub>Hydrophilic</sub> /nm<sup>2</sup> | 23.1±1.9 | 22.3±1.6 | 22.1±1.5 | 21.9±1.7 | 24.1±1.6 | n/a |
| Dimer SASA<sub>Hydrophilic</sub> /nm<sup>2</sup> | 37.2±3.8 | 34.8±2.9 | 36.3±3.3 | 34.6±3.2 | 38.7±3.3 | n/a |
| ΔSASA<sub>Hydrophobic</sub> /nm<sup>2</sup> | -8.4±4.0 | -9.8±2.6 | -8.2±3.4 | -8.9±3.4 | -10.2±3.5 | n/a |
| ΔSASA<sub>Hydrophilic</sub> /nm<sup>2</sup> | -9.0±4.0 | -9.9±3.1 | -8.0±3.4 | -9.3±3.7 | -9.5±3.6 | n/a |
| Inter-chain BB-BB | 19±13 | 20±9 | 10±9 | 14±10 | 23±13 | n/a |
| Intra-chain BB-BB | 21±11 | 23±10 | 12±7 | 16±9 | 21±12 | n/a |
| Inter-chain SC-SC | 59±26 | 70±17 | 56±22 | 65±21 | 70±22 | n/a |
| Intra-chain SC-SC | 64±18 | 73±16 | 54±15 | 65±17 | 59±17 | n/a |
| Total H-bonds | 42.4±6.6 | 47.9±6.0 | 51.1±7.1 | 38.4±6.2 | 33.3±6.0 | n/a |
| Intra-chain A H-bonds | 16.7±4.3 | 19.1±4.7 | 22.7±4.9 | 15.6±4.3 | 11.6±4.1 | n/a |
| Intra-chain B H-bonds | 16.9±4.2 | 19.4±4.4 | 22.5±4.8 | 15.3±4.0 | 11.2±4.1 | n/a |
| Inter-chain H-bonds | 8.8±5.4 | 9.5±4.2 | 5.9±4.2 | 7.5±4.3 | 10.5±5.4 | n/a |
| $\Delta E_{\text{binding}}$ / kJ·mol<sup>-1</sup> | 90.9±4.1 | 19.0±3.4 | 69.5±4.0 | 150.3±4.4 | -93.6±6.3 | n/a |
| $\Delta G_{\text{polar}} / \text{kJ}\cdot\text{mol}^{-1}$ | 561.0±4.6 | 619.9±4.0 | 527.8±4.1 | 580.7±4.5 | 543.7±5.5 | n/a |
| $\Delta G_{\text{non-polar}} / \text{kJ}\cdot\text{mol}^{-1}$ | -40.9±0.3 | -46.3±0.2 | -37.6±0.2 | -43.8±0.2 | -47.3±0.3 | n/a |

<sup>a</sup> Levene's Test p-values, and ANOVA F-statistic and p-values for ANOVA are listed in Table S1.
<sup>b</sup> Welch's ANOVA p-values were < 0.0001 except for c where Levene's p-value was 0.1.
<sup>c</sup> Indicates the use of standard ANOVA based on the assumption for homogeneity of variance being met as demonstrated by Levene's Test.

No homogeneity exists in the variances for all calculated values across all force fields with Levene’s test p-values <0.0001. The exception is the individual chain SASA_Hydrophobic_ with a p-value = 0.1. These p-values indicate that further extension of the simulations would be statistically unlikely to change the level of homogeneity in the variance. The results of Welch’s ANOVA indicate significant differences between the results obtained with the force fields (and when available experimental data and previously published dimer simulations)^23,26^ with p-values <0.0001. Similar to the conclusion drawn from Levene’s test these p-values indicate that further extension of the simulations would be statistically unlikely to change the results. Differences in P_2_, R_g_, CCS, fractions of populated secondary structure (β-sheet, α-helix, and turn/coil/bend), SASA of the individual chains and dimer, ISAs, SASA_Hydrophobic_ of the individual chains and dimer, SASA_Hydrophilic_ of the individual chains and dimer, the change in hydrophobic and hydrophilic surface areas with dimerization, and the number of intra- and inter-chain BB-BB and SC-SC contacts are inherent properties of the force fields as utilized for this system.

Welch’s ANOVA is a statistically robust method of determining if there are statistically significant differences between the means of the calculated values obtained by different force fields (and experimental data when available).^92-97^ The analysis, however, cannot determine which differences between means, if present, are statistically significant. To determine the nature of these differences, a *post-hoc* Games-Howell test was used (Table 2).^100^ The *post-hoc* analysis demonstrates that there are no systematic similarities between any of the force fields and statistically significant similarities occur on sporadic basis that are independent of the calculated properties.

**Table 2.** Non-statistically significant comparisons between force fields as determined by Games-Howell *post-hoc* analysis, all other comparisons are considered statistically significant.

| Calculated Property | Compared Force Fields | | Mean Difference ± Standard Deviation |
| --- | --- | --- | --- |
| **α-Helix /%** | AMBER-ILDN* | CHARMM22* | 0.6±0.18 |
| **Turn, Coil, Bend /%** | AMBER-ILDN | CHARMM22* | 0.0±0.19 |
| **Individual Chain SASA<sub>Hydrophobic</sub> /nm<sup>2</sup>** | AMBER-NMR | AMBER-ILDN* | 0.0±0.08 |
| **Dimer SASA<sub>Hydrophobic</sub> /nm<sup>2</sup>** | AMBER-NMR | CHARMM22* | 0.1±0.13 |
| **Dimer SASA<sub>Hydrophilic</sub> /nm<sup>2</sup>** | AMBER-ILDN* | CHARMM22* | 0.2±0.11 |
| **ΔSASA<sub>Hydrophobic</sub> /nm<sup>2</sup>** | AMBER-NMR | AMBER-ILDN | 0.2±0.12 |
| **ΔSASA<sub>Hydrophilic</sub> /nm<sup>2</sup>** | AMBER-ILDN | CHARMM22* | 0.3±0.12 |
| **ΔSASA<sub>Hydrophilic</sub> /nm<sup>2</sup>** | CHARMM36 | CHARMM22* | 0.2±0.12 |

For all simulations and force fields, <*P*_*2*_> and <R_g_> for the individual chain and dimer remain stable throughout the production phase. Values of <*P*_*2*_> indicate that the sampled conformers for all simulations represent fluctuating soluble nematic droplets with relatively high fraction of β-sheet content.^57,62^ *Post-hoc* analysis showed the largest difference between AMBER-NMR and CHARMM36, and the smallest difference between CHARMM22* and AMBER-NMR. All comparisons showed statistically significant differences. Different force fields results in statistically significant differences, although the effects are minor. The effect of force field selection on <R_g_> and CCS was significant for all simulations indicating ensembles of conformations that are slightly larger than measured by IM-MS.^105,106^ The trend in values of <R_g_> and CCS are AMBER-ILDN > CHARMM36 > AMBER-NMR > CHARMM22* > AMBER-ILDN*. The greatest degree of variability occurs within AMBER-NMR and AMBER-ILDN for <R_g_> and AMBER-NMR, AMBER-ILDN, and CHARMM22* for CCS. These findings are likely secondary to differences in force field and solvent model parameters^107-109^

### Secondary Structure and Cα-Backbone Stability

There are marked differences in the fractions of populated DSSP calculated secondary structure (helix, β-sheet, and turn/coil/bend) for each of the simulations when compared to experimental results (Table 1 and Table S2). The AMBER-NMR results show significant oversampling of α-helical content compared to CD, while CHARMM36^26^ tended to under sample this conformation by a small but significant amount. There was a statistically significant similarity in α-helical sampling between AMBER-ILDN* and CHARMM22*. All force fields tended to over sample the fraction of turn/coil/bend compared to CD^104,110^ with the exception of AMBER-NMR. It should be noted, however, that this force-field was heavily weighted by α-helical sampling which accounts for this result. The AMBER-ILDN and CHARMM22* were statistically similar in their degree of global turn/coil/bend conformational sampling. None of the force fields were able to accurately sample the fraction of β-sheet content with the best performing force field being CHARMM36 followed by AMBER-ILDN > AMBER-ILDN* > CHARMM22* > AMBER-NMR.

The fraction of sampled secondary structure and degree of chain flexibility as measured by the Cα-atom RMSF of the individual residues as a function of residue number of Chain A and Chain B for each of the force fields are shown in Figure 4. Similar results were obtained if the Cα-atoms of the dimer system are overlaid on the average conformation of the dimer, with higher values (avg. 0.2 nm/residue).

**Figure 4.**
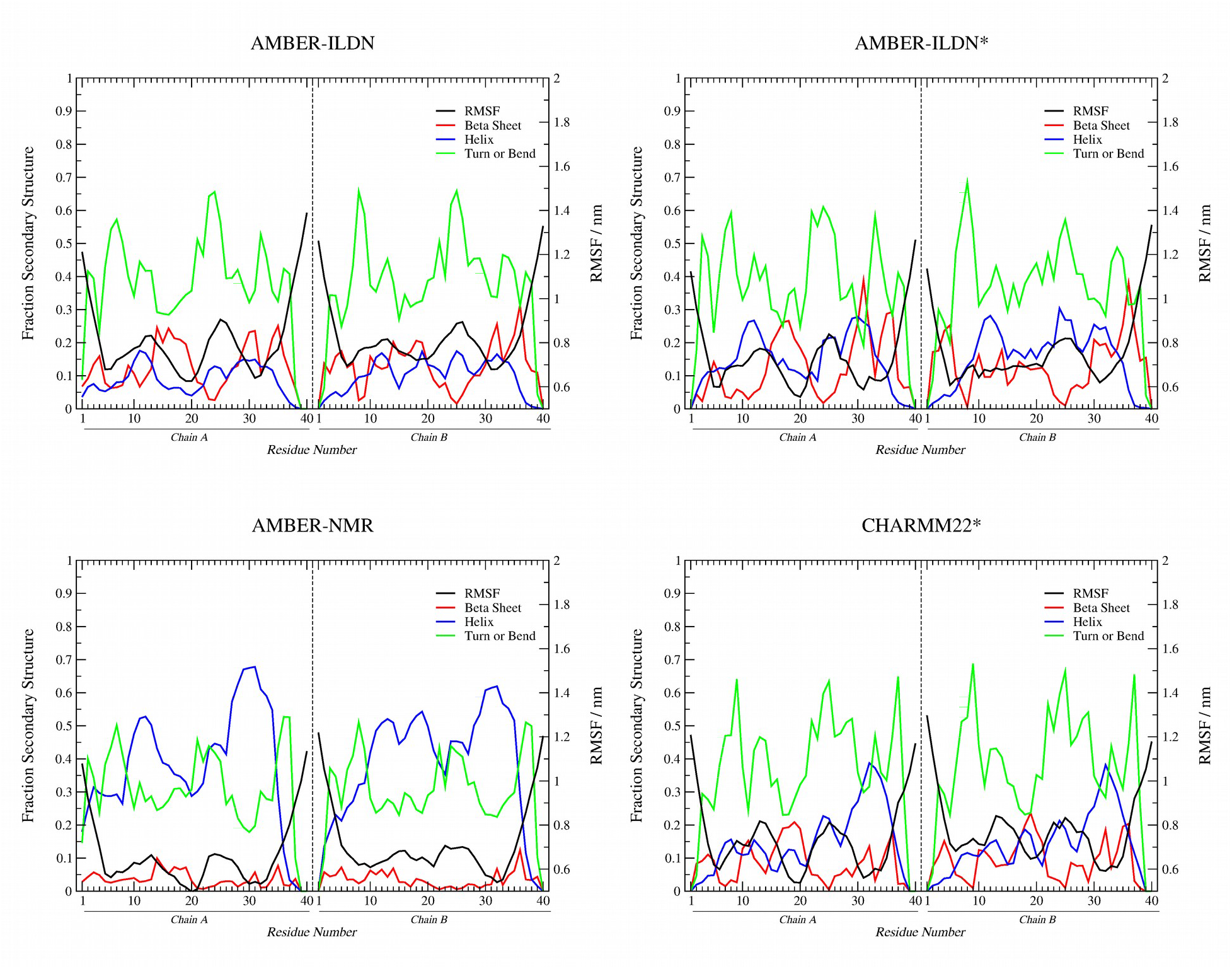
Backbone RMSF for each monomer chain overlaid with its own average structure for AMBER-ILDN, AMBER-ILDN*, AMBER-NMR, and CHARMM22* force fields. The RMSF data compared to a plot of the fraction of sampled secondary structures as a function of residue number. RMSF, black; helix, blue; β-sheet, red; β-turn/bend, green.

### Solvent Exposed Surface Area

The trend in values of SASA (Table 1) for the different force fields for the monomer is CHARMM36 > AMBER-ILDN > CHARMM22* > AMBER-ILDN* > AMBER-NMR and the dimer is CHARMM36 > AMBER-ILDN > AMBER-NMR > CHARMM22* > AMBER-ILDN* in rough agreement with the results from R_g_ and CCS for the size of the dimer and the individual chains. CHARMM36 also demonstrates the largest ISA followed by AMBER-ILDN* and CHARMM22* while AMBER-ILDN and AMBER-NMR have the least. For hydrophobic and hydrophilic surface areas, CHARMM36 and AMBER-ILDN are again the largest followed by CHARMM22*, AMBER-NMR, and AMBER-ILDN*. The ratios of hydrophobic to hydrophilic SASA for the dimer are: AMBER-ILDN*, 1.54; AMBER-NMR, 1.49; CHARMM36, 1.47; CHARMM22*, 1.42; and AMBER-ILDN, 1.39. There are statistically significant similarities between the individual chain SASA_Hydrophobic_ values of AMBER-NMR and AMBER-ILDN*, the dimer SASA_Hydrophobic_ values of AMBER-NMR and CHARMM22*, and the dimer SASA_Hydrophilic_ values of AMBER-ILDN* and CHARMM22*. With dimerization, CHARMM36 demonstrates^26^ a slightly larger decrease in the amount hydrophobic solvent exposed surface compared to hydrophilic solvent exposed surface area while AMBER-ILDN and CHARMM22* show the opposite relationship. There are statistically significant similarities in the ∆SASA_Hydrophobic_ between AMBER-NMR and AMBER-ILDN, ∆SASA_Hydrophilic_ between AMBER-ILDN and CHARMM22* and CHARMM36 and CHARMM22*.

The rSASA and rISA for different force fields are shown in Figure 5. For AMBER-ILDN force field, the average rSASA of the dimer is 0.49±0.11 with maximal values occurring at the N- and C-terminal residues with a small peak from residues 21-28 of both chains. The average rISA is 0.13±0.03 with a maximal value of 0.23 and minimal value of 0.05. The rISA of the dimer is relatively flat without a significant degree of solvent shielding or collapse of the hydrophobic core regions. For AMBER-ILDN* force field, the average rSASA of the dimer is 0.44±0.12 with maximal values occurring at the N- and C-terminal residues with a small peak from residues 21-28 of both chains. The average rISA is 0.15±0.04 with a maximal value of 0.27 and minimal value of 0.05. The rISA of the dimer is again relatively flat with a minor degree of solvent shielding that is more prominent at the N-terminal regions of both chains from residues 31-38. For AMBER-NMR force field, the average rSASA of the dimer is 0.46±0.12 with maximal values occurring at the N- and C-terminal residues with a small nadir from residues 28-33 of both chains. The average rISA is 0.12±0.03 with a maximal value of 0.22 and minimal value of 0.05. The rISA of the dimer is again relatively flat. For CHARMM22*, the average rSASA of the dimer is 0.46±0.12 with maximal values occurring at the N- and C-terminal residues with two small nadirs from residues 17-21 and 30-35 of both chains. The average rISA is 0.14±0.04 with a maximal value of 0.25 and minimal value of 0.07. The rISA of the dimer is again relatively flat despite the relative shielding of residues 17-21 and 30-35 ***Intra- and Inter-chain Interactions.***

**Figure 5.**
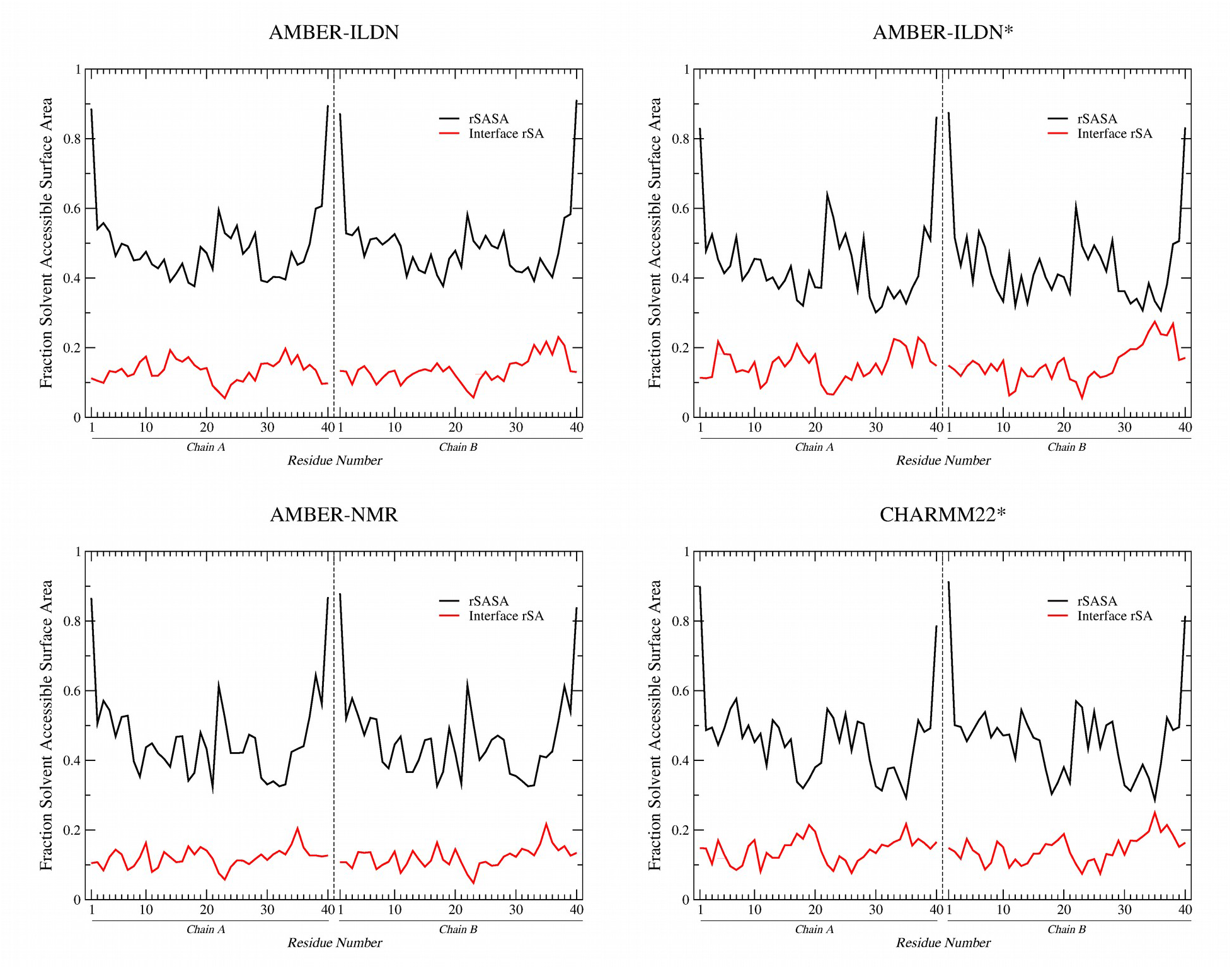
Relative solvent accessible surface area (rSASA) for the Aβ(1-40) dimer as a function of residue number compared to the relative surface area of the interaction (rISA) between the Chain A and Chain B for AMBER-ILDN, AMBER-ILDN*, AMBER-NMR, and CHARMM22* force fields. rSASA, black; rISA, red.

### Residue Contacts

The mean number of inter-chain BB-BB contacts (Table 1.) for the different force fields follows the trend of CHARMM36 > AMBER-ILDN* > AMBER-ILDN > CHARMM22* > AMBER-NMR while intra-chain BB-BB contacts follows the trend AMBER-ILDN* > CHARMM36 ~ AMBER-ILDN > CHARMM22* > AMBER-NMR. The CHARMM36 and AMBER-ILDN* force fields also have the largest number of inter-chain SC-SC contacts followed by CHARMM22* > AMBER-ILDN > AMBER-NMR force fields. The intra-chain contact trend is AMBER-ILDN* > CHARMM22* ~ AMBER-ILDN > CHARMM36 > AMBER-NMR force fields. The interactions within the CHARMM36^26^ force field show a general trend to being dominated by inter-chain interactions, either BB-BB or SC-SC while the AMBER-NMR force field demonstrated the lowest number of dimer contacts and interactions. There were no statistically significant similarities between any of the force fields for the number of intra-chain BB-BB, inter-chain BB-BB, intra-chain SC-SC, or inter-chain SC-SC contacts. The inter- and intra-chain residue contact probability maps for backbone-backbone (BB-BB) and side chain-side chain (SC-SC) contacts for each force-field are shown in Figure 6.

**Figure 6.**
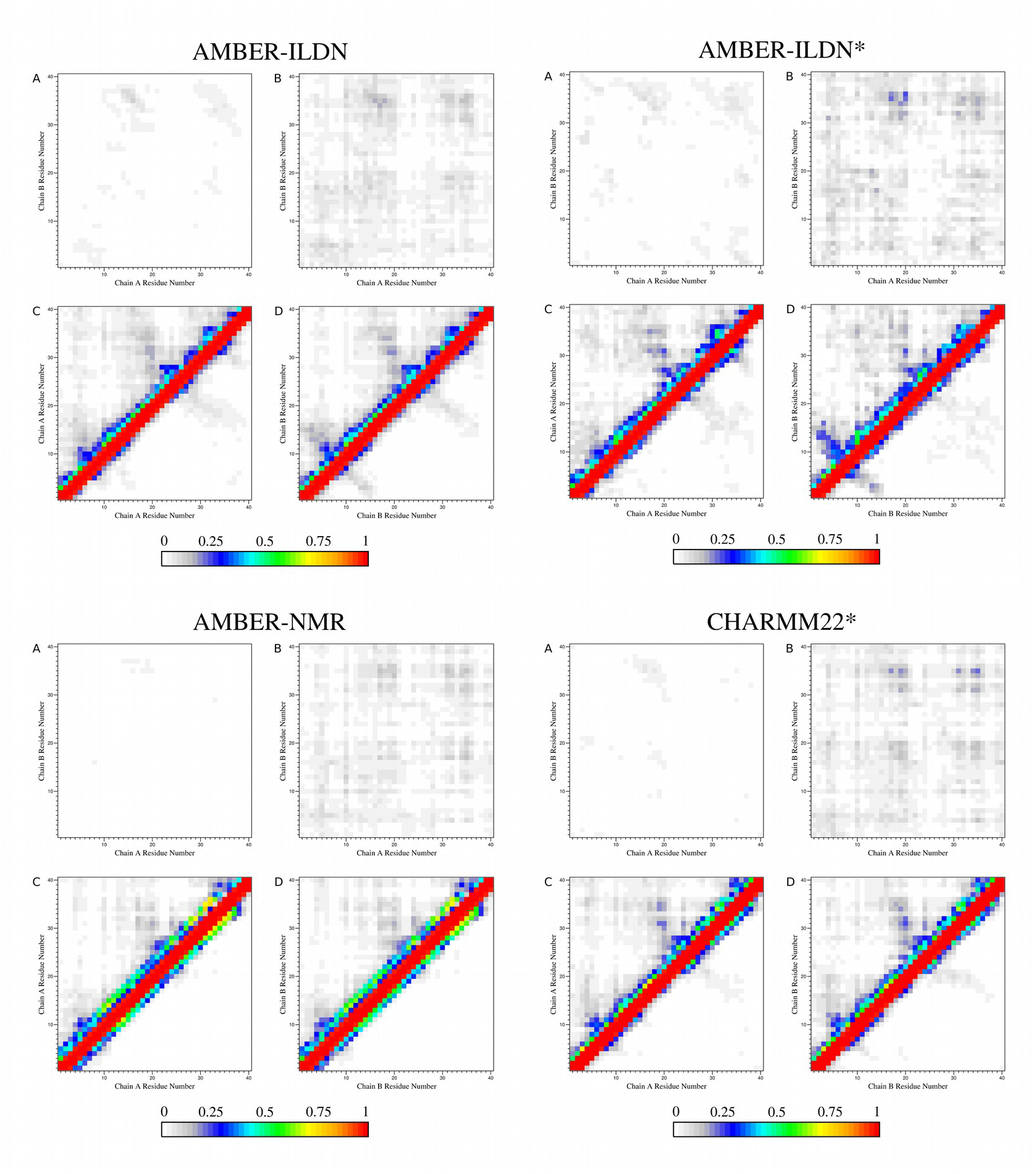
Inter- and intra-chain residue contact probability maps for AMBER-ILDN, AMBER-ILDN*, AMBER-NMR, and CHARMM22* force fields. **A**, backbone-backbone (BB-BB) contacts of chain A and chain B; **B**, side chain-side chain (SC-SC) contacts of chain A and chain B; **C**, BB-BB and SC-SC contacts of chain A with itself; **D**, BB-BB and SC-SC contacts of chain B with itself. For the BB-BB and SC-SC intra-chain contacts (**C**) and (**D**) the BB-BB contacts are shown above the diagonal while SC-SC contacts are shown below the diagonal. Color scale bars represent the relative probabilities of residue contact.

### Hydrogen bonds

The mean number of intra- and inter-chain H-bonds (Table 1) follow the trend AMBER-NMR > AMBER-ILDN* > AMBER-ILDN > CHARMM22* > CHARMM36. There were no statistically significant similarities between any of the force fields for the number of intra-chain, or inter-chain hydrogen bonds. The normalized fractions of per residue occupancy for the intra- and inter chain hydrogen bonds of each force field are shown in Figure 7. The per residue occupancy for all force fields are very low < 0.15 for the intra-chain interactions and < 0.10 for the inter-chain interactions.

**Figure 7.**
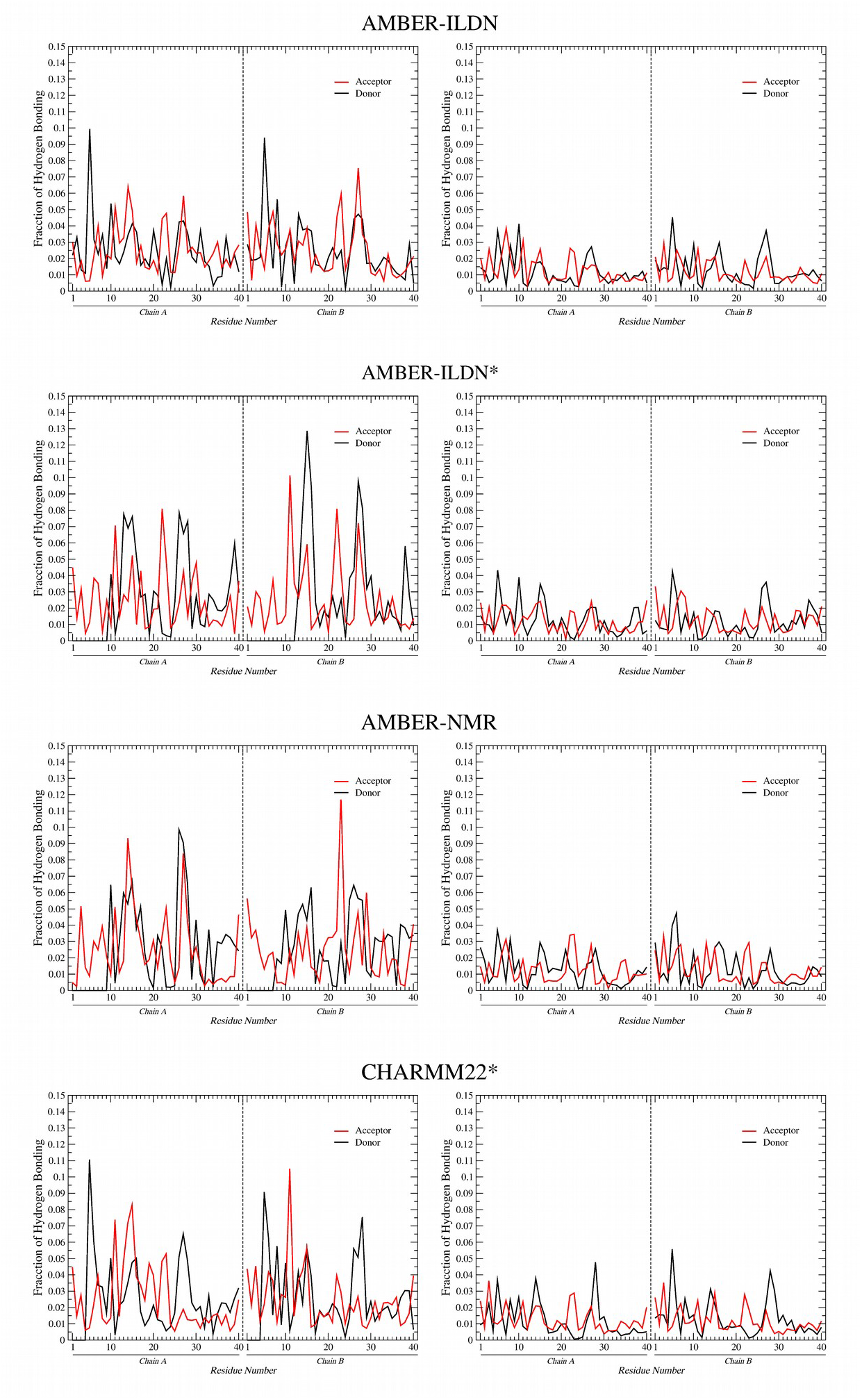
Fraction of populated H-bonds normalized to the total number of H-bonds observed during the simulation plotted as a function of residue number for AMBER-ILDN, AMBER-ILDN*, AMBER-NMR, and CHARMM22* force fields. The intra-chain donor acceptor residues are on the right panels and the inter-chain donor acceptor residues are on the left panels.

### Binding energies

The binding energies (∆E_binding_) of chain A to chain B for the simulations at 310 K are given in Table 3. Of the surveyed force fields, only CHARMM36 (ΔE_binding_ = −93.560 ± 6 kJ•mol-^1^)^26^ demonstrates an energetically favorable ∆E_binding_ for the interaction between the individual chains. The components of ∆E_binding_ (Table 3) demonstrate that the ∆G_polar_ desolvation energy (∆G_polar,CHARMM36_ = 543.671 ± 5.531^26^) is the major destabilizing factor for the dimer. There are neither discernable trends nor statistically significant similarities in ∆E_vdw_, ∆E_elec_, ∆G_polar_, and/or ∆G_non-polar_ values that explains differences between any of the force fields.

**Table 3.**
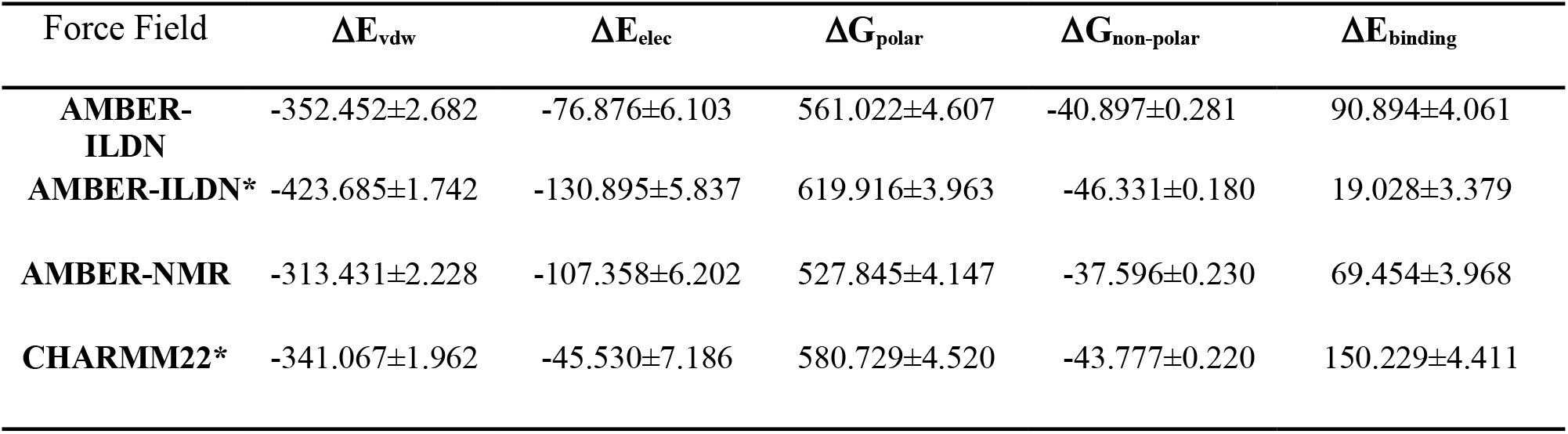
Binding energy (ΔE_binding_) ± standard error and its components for binding monomers of the Aβ(1-40). All units are in kJ•mol^−1^.

Per residue contributions to ∆E_binding_ are shown in Figure 8. Results for all simulations and force fields are essentially superimposable with only minor variations. The acidic residues, Asp and Glu and the terminal Val have large positive contributions, whereas, the two basic residues Arg and Lys have large negative contributions to the binding energy. The side chain hydration of the Arg, Asp, Glu and Lys residues were evaluated by integrating the radial distribution function within 0.5 nm distance of the sidechain’s terminal moiety (Table 4). The integral of this function is equal to the probability (P_*g(r)*_) of finding water molecules within a defined radius. Examination of the data reveals subtle difference between the force fields that range from 0.01 to 0.06 on a per residues basis. For the acidic sidechains of Asp and Glu the probabilities range from 0.23-0.30. Whereas, the basic side chains of Arg and Lys probabilities range from 0.15-0.23. These probabilities indicate that it is easier to desolvate the basic residues than the acidic residues and this is in agreement with the data of White and colleagues who showed that the order of free energy of desolvation is Asp ~ Glu > Lys > Arg.^111^

**Figure 8.**
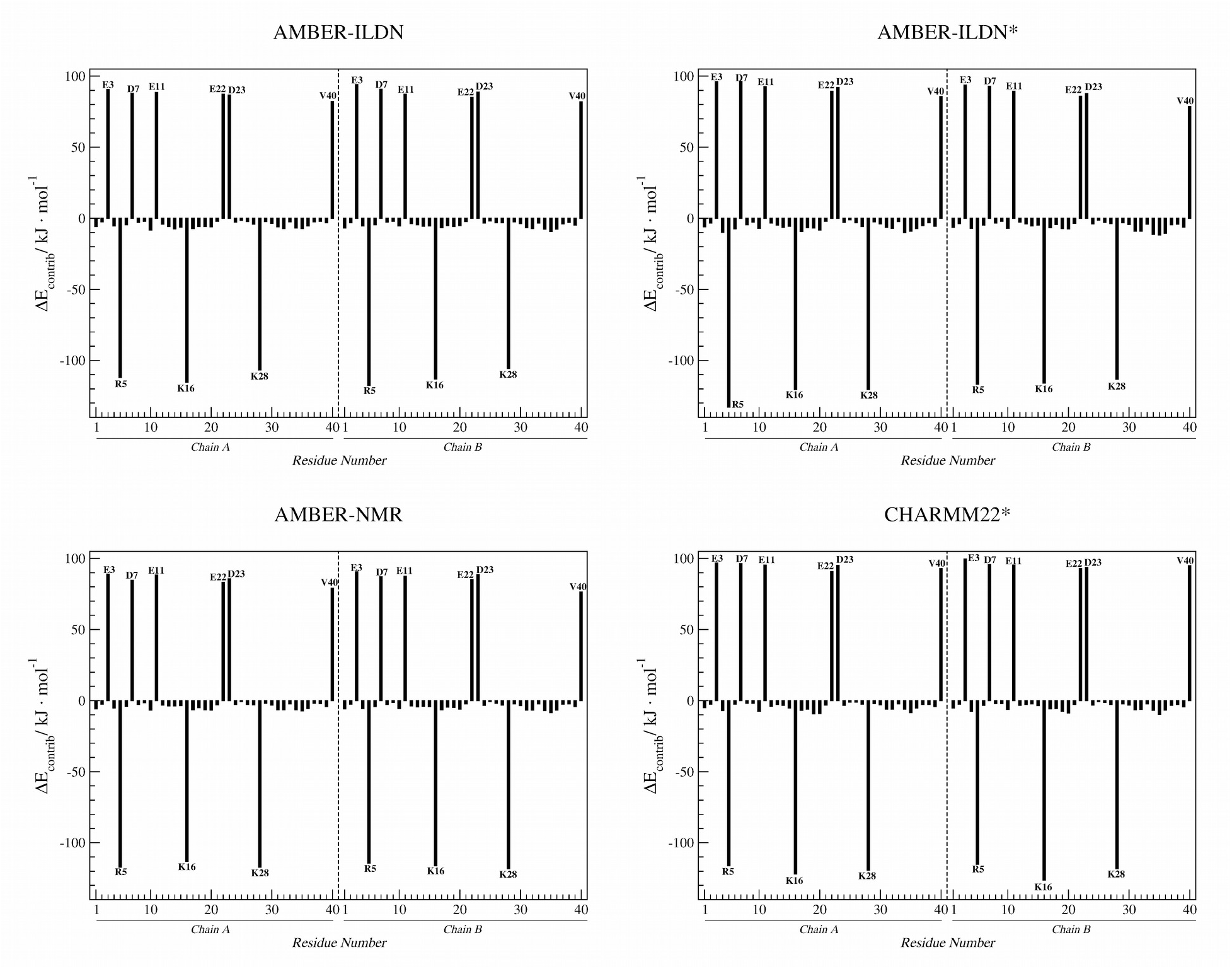
Contribution of residues to the binding energy of the Aβ(1-40) dimer for AMBER-ILDN, AMBER-ILDN*, AMBER-NMR, and CHARMM22* force fields. Residues with large energy contributions are indicated by one letter code.

**Table 4.** Probability (P_*g(r)*_) of finding water molecules within 0.5 nm distance of side chain atoms of acidic and basic residues.

| Chain | Residue | AMBER-ILDN | AMBER-ILDN* | AMBER-NMR | CHARMM22* |
| --- | --- | --- | --- | --- | --- |
| **A** | Glu<sup>3</sup> | 0.25 | 0.24 | 0.24 | 0.23 |
|  | Arg<sup>5</sup> | 0.16 | 0.15 | 0.16 | 0.16 |
|  | Asp<sup>7</sup> | 0.23 | 0.23 | 0.24 | 0.24 |
|  | Glu<sup>11</sup> | 0.25 | 0.25 | 0.26 | 0.25 |
|  | Lys<sup>16</sup> | 0.21 | 0.22 | 0.23 | 0.22 |
|  | Glu<sup>22</sup> | 0.29 | 0.3 | 0.29 | 0.26 |
|  | Asp<sup>23</sup> | 0.26 | 0.25 | 0.24 | 0.25 |
|  | Lys<sup>28</sup> | 0.24 | 0.23 | 0.23 | 0.22 |
| **B** | Glu<sup>3</sup> | 0.25 | 0.22 | 0.25 | 0.23 |
|  | Arg<sup>5</sup> | 0.15 | 0.15 | 0.16 | 0.16 |
|  | Asp<sup>7</sup> | 0.23 | 0.22 | 0.23 | 0.24 |
|  | Glu<sup>11</sup> | 0.28 | 0.26 | 0.26 | 0.24 |
|  | Lys<sup>16</sup> | 0.22 | 0.21 | 0.23 | 0.21 |
|  | Glu<sup>22</sup> | 0.29 | 0.28 | 0.29 | 0.27 |
|  | Asp<sup>23</sup> | 0.25 | 0.24 | 0.24 | 0.25 |
|  | Lys<sup>28</sup> | 0.24 | 0.22 | 0.22 | 0.23 |

The DCCMs shown in Figure 9, demonstrate that for the AMBER-ILDN, AMBER-ILDN*, and AMBER-NMR force fields, there are strong correlated intra-chain motions but strongly anti-correlated inter-chain motions for the dimer. For the CHARMM22* force field, there is a weakly positive correlation between the central hydrophilic region corresponding to the loop region with the N-terminal hydrophilic regions. Only the CHARMM36 force field demonstrates strongly correlated motions between the two chains which are consistent with favorable interactions and the strong binding energy.

**Figure 9.**
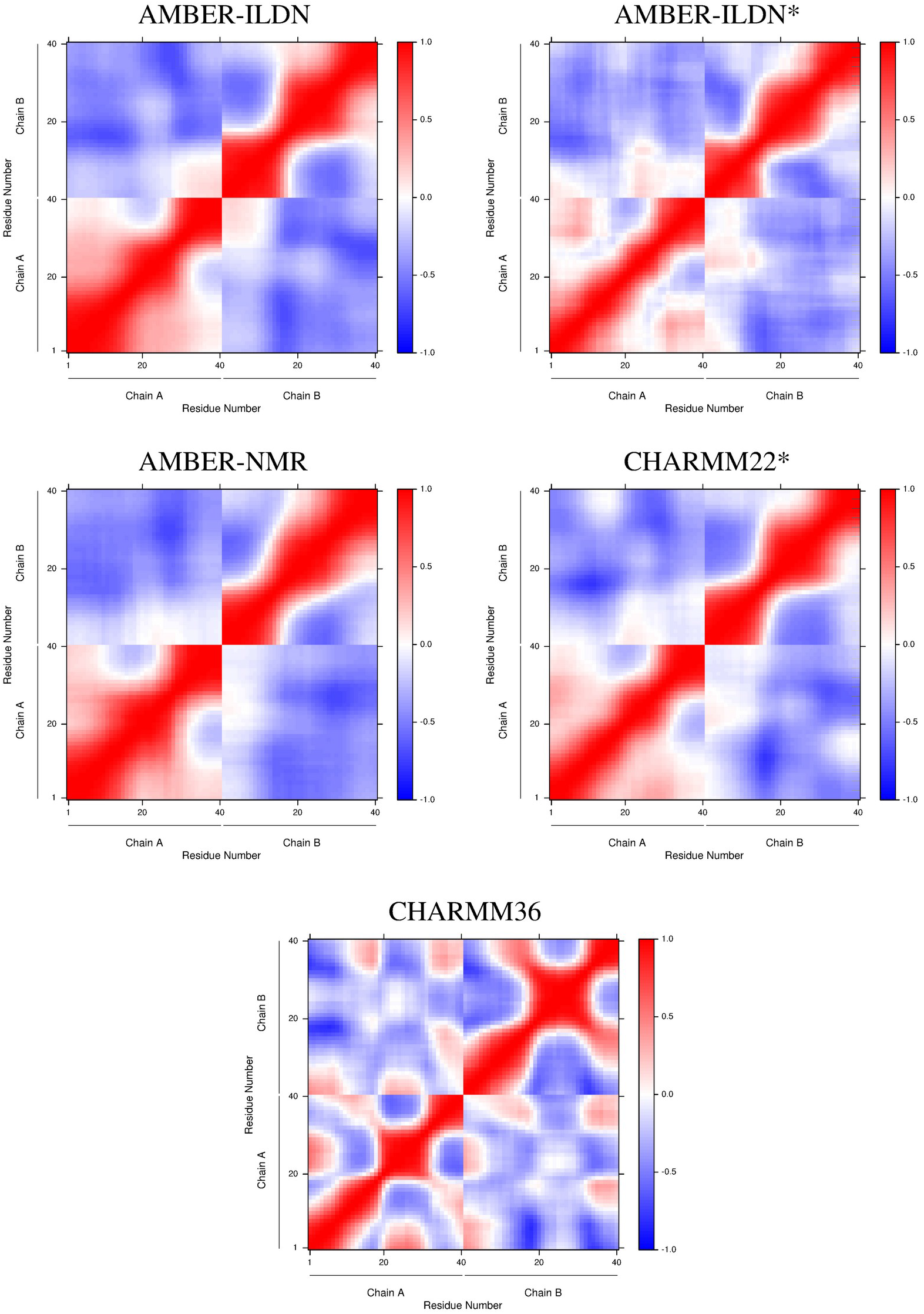
Dynamics cross-correlation matrices (DCCM) for AMBER-ILDN, AMBER-ILDN*, AMBER-NMR, CHARMM22*, and CHARMM36 force fields. Values range from −1 (complete anticorrelation) to +1 (complete correlation).

### Dihedral Principal Component Analysis

The lowest energy conformations were determined by projecting each trajectory onto the first three dihedral principal components as show in Figure 10. Free energy landscapes for all four force fields have shallow minima with a maximum of 5.7 kJ•mol^−1^ separating the lowest and highest energy groups. The separation between highest and lowest energy groups were: 5.0 kJ•mol^−1^, 5.7 kJ•mol^−1^, 4.6 kJ•mol^−1^, and 4.1 kJ•mol^−1^, respectively, AMBER-ILDN, AMBER-ILDN*, AMBER-NMR, and CHARMM22* with a previously published value of 4.1 kJ•mol^−1^ for CHARMM36.^26^ Secondary structure analysis was performed using the DSSP method (structures in the figure may have small differences since YASARA uses the STRIDE algorithm).^112^

**Figure 10.**
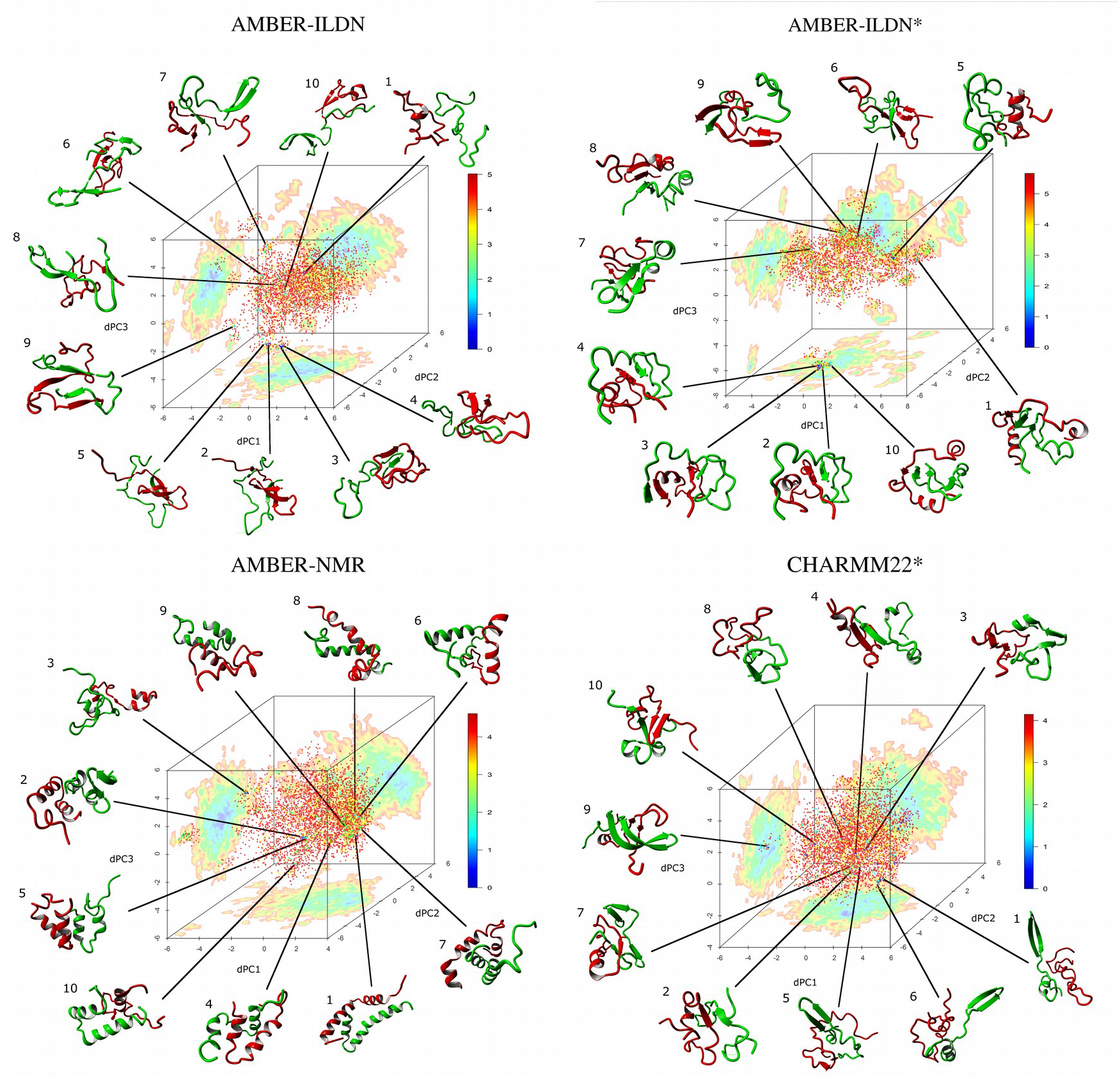
Free energy landscapes of the dimer (kJ•mol^−1^) as a function of the first three components of the dihedral principle component analysis (dPC1, dPC2, dPC3) for AMBER-ILDN, AMBER-ILDN*, AMBER-NMR, and CHARMM22*. The associated two dimensional projections of the free energy landscape are also displayed. The ten lowest energy conformations shown and numbered in order from lowest to highest with chain A in red and chain B in green. Arrows represent the β-sheet conformation. The conformations have been rotated to optimally display the secondary structural elements and dimer interactions.

The lowest energy representative structure for AMBER-ILDN force field had a dimer SASA of 72.31 nm^2^ with an ISA of 10.10 nm^2^. The dimer was less compact (R_g_ = 1.43 nm) than chain A (R_g_ = 1.08 nm) and chain B (R_g_ = 1.20 nm). A total of 5 salt bridges were present: chain A Glu^3^-Arg^5^, chain A Glu^11^-chain B His^13^, and chain B Asp^7^-Arg^5^-Glu^11^-His^13^. There was a single intra-chain aromatic-aromatic (Ar-Ar) interaction between chain A His^13^ and chain B His^14^. A single β-bridge was found between residue 12 of chain A and residue 13 of chain B, and a single α-helix was found in chain A at residues 29-32.

The lowest energy representative structure for AMBER-ILDN* force field had a dimer SASA of 56.90 nm^2^ with an ISA of 24.60 nm^2^. The dimer was more compact (R_g_ = 1.26 nm) than chain A (R_g_ = 1.42 nm) and less compact than chain B (R_g_ = 1.02 nm). A total of 5 salt bridges were present: chain A His^6^-Asp^7^, chain A His^14^-chain B Asp^7^, chain A Arg^5^-chain B Asp^23^, chain B Glu^3^-Arg^5^ and Asp^7^-Lys^16^. The secondary structure was 24% helix with α-helices in chain A at residues 3 to 6 and 28-35, in chain B at residues 16 to 19. A single 3 _10_ helix was found at residues 20 to 22 in chain A. Two intra-chain anti-parallel β-bridges were found between residues 12-29 and 6-38 of chain B. An inter-chain parallel β-sheet forms between residues 18-19 of chain A and 38-39 of chain B.

The lowest energy representative structure for AMBER-NMR force field had a dimer SASA of 58.59 nm^2^ with an ISA of 16.27 nm^2^. The dimer was less compact (R_g_ = 1.45 nm) than chain A (R_g_ = 1.44 nm) and chain B (R_g_ = 1.29 nm). A total of 2 salt bridges were present: chain A Arg^5^-Asp^7^, chain A Glu^3^-chain B Arg^5^. There was 1 intra-chain and 2 inter-chain Ar-Ar interactions between chains A Phe^19^-Phe^20^, chain A Tyr^10^-chain B His^13^, and chain A His^13^-chain B Tyr^10^. The secondary structure was 66% α-helix with helices running from residues 6-15 and 21-35 in chain A and 4-17, 19-25, and 30-36 in chain B.

The lowest energy representative structure for CHARMM22* force field had a dimer SASA of 54.28 nm^2^ with an ISA of 16.27 nm^2^. The dimer was less compact (R_g_ = 1.42 nm) than chain A (R_g_ = 1.04 nm) and more compact than chain B (R_g_ = 1.45 nm). A total of 4 salt bridges were present: chain A Asp^1^-Arg^5^ and Asp^23^-Lys^28^, chain B Arg^5^-Glu^3^-Lys^16^. There was a single intra-chain Ar-Ar interaction between Phe^19^-Phe^20^. Chain A contained 3_10_ helices at residues 17-21 and 27-30 and an α-helix at residues 31-34. Chain B contained an α-helix at residues 28-34 and an antiparallel β-sheet formed between residues 2-7 and 10-15.

### Essential Subspace Analysis

A comparison of the sampled essential subspace was performed by calculating the RMSIP and nRMSIP for each force field pair as shown in Figure 11. The use of a normalized value for the RMSIP should in theory account for difference in values secondary to the effects of autocorrelation and sampling error.^89^ The effects of this correction are seen particularly with respect to the differences between AMBER-ILDN* and CHARMM36 where the RMSIP wound suggest that the fraction of shared essential subspace is lower at 0.764 than the corrected value of 0.900. Despite differences in the lowest energy conformations (Figure 10) all force fields have a significant degree of similarity with regards to essential subspace sampling with all normalized values within 0.100 of unity.

**Figure 11.**
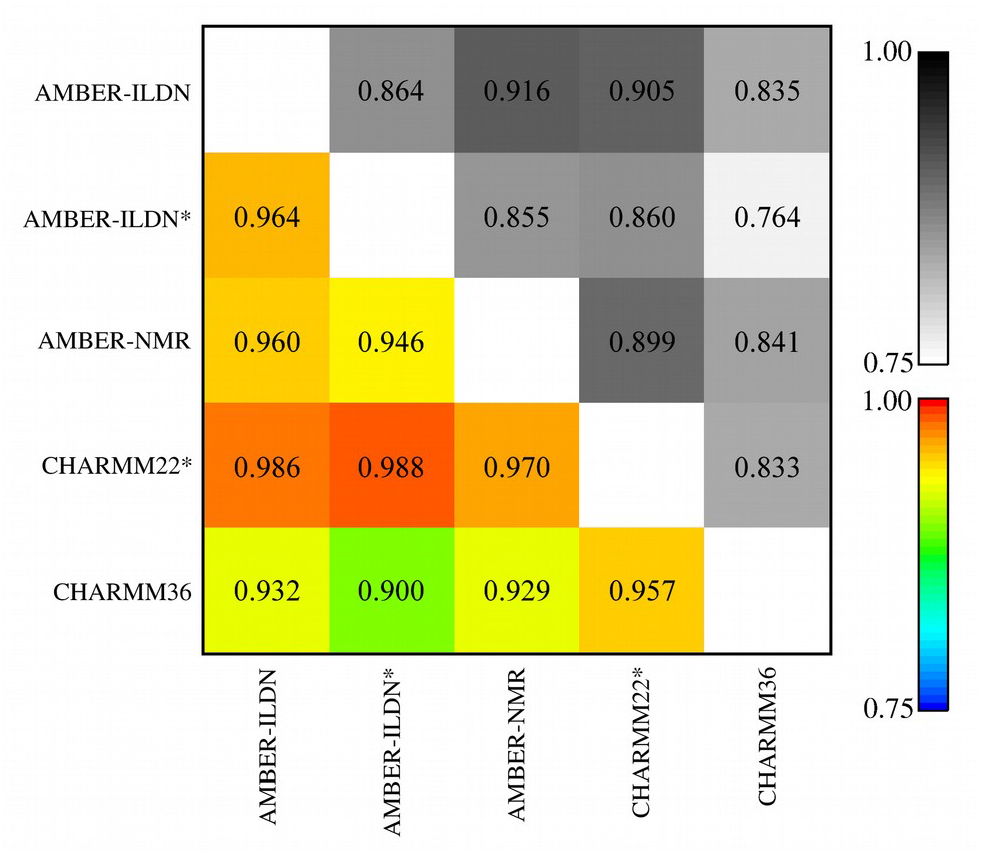
The root mean square inner product (RMSIP) and normalized RMSIP (nRMSIP) overlap matrix of AMBER-ILDN, AMBER-ILDN*, AMBER-NMR, CHARMM22*, and CHARMM36 demonstrating the comparison of essential subspace sampling similarity between force fields. The RMSIP is shown on the top right in greyscale and the nRMSIP on the bottom left in color. By definition, the diagonal values are equal to 1.0.

Differences in essential subspace sampling a three dimensional overlay of the first three dPCs with the two dimensional projections of dPC1•dPC2, dPC1•dPC3, and dPC2•dPC3 are shown in Figure 12.^89^ The vast majority of the sampled subspace are tightly overlaid for all force fields with only minor outlying differences. There were no statistically significant differences comparing the sampled dihedral essential subspace for dPC1 and dPC3 (Table 5). There was a statistically significant difference in dPC2, but it was not demonstrated in the subsequent Games-Howell analysis and is therefore most likely artefactual.

**Figure 12.**
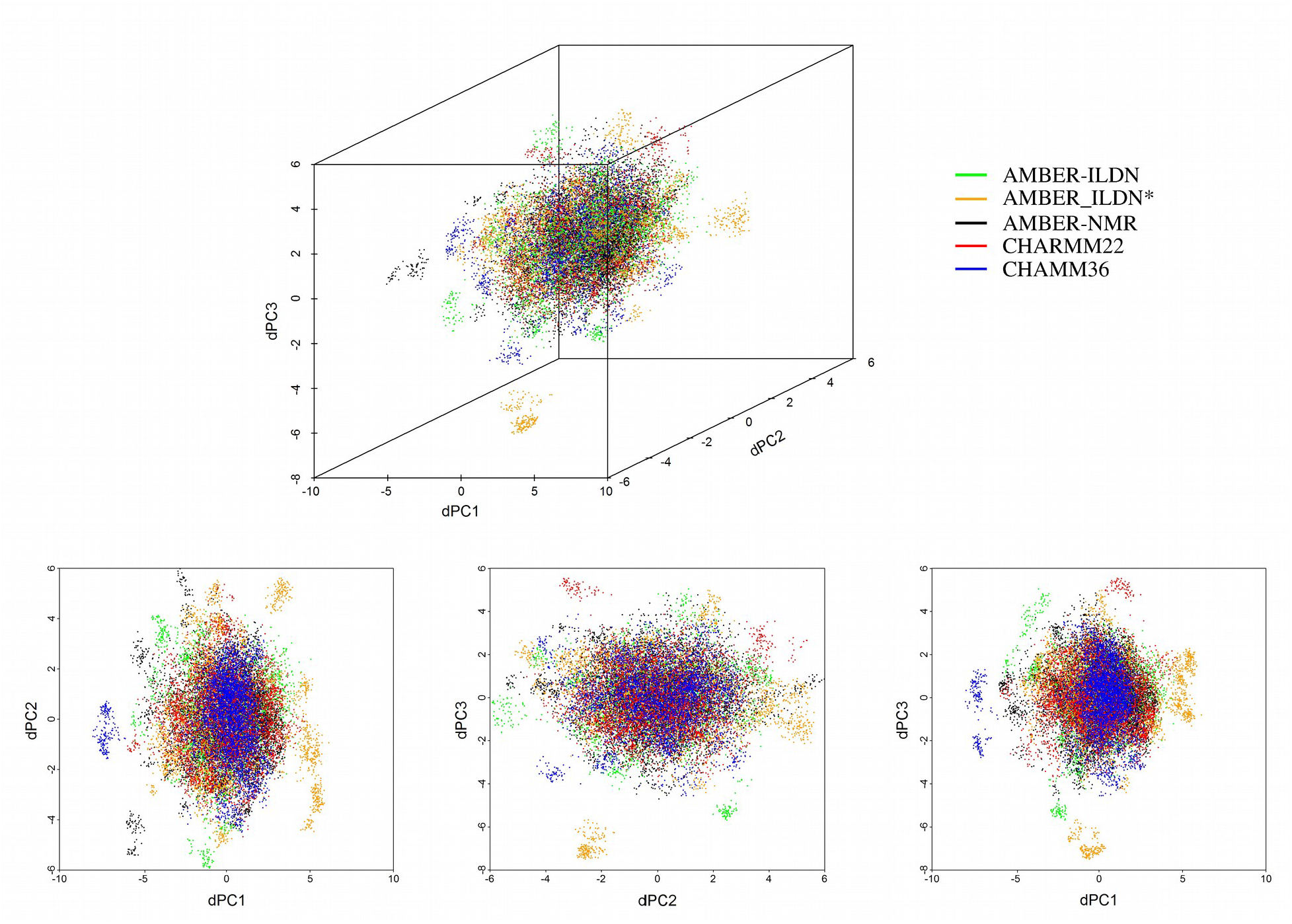
Three dimensional overlay of the first three components of the dihedral principal component analysis (dPC) with the two dimensional projections of dPC1•dPC2, dPC1•dPC3, and dPC2•dPC3 for AMBER-ILDN, AMBER-ILDN*, AMBER-NMR, CHARMM22*, and CHARMM36.

**Table 5.** ANOVA for dihedral principal components (dPC1-3).

| dPC | Mean ± Standard Deviation | | | | | Levene's | Welch's |
| --- | --- | --- | --- | --- | --- | --- | --- |
|  | AMBER-ILDN | AMBER-ILDN* | AMBER-NMR | CHARMM22* | CHARMM3<br>6 | p-value | ANOVA<sup>a</sup><br>p-value |
| dPC1 | 0.00 | 0.00 | -0.08 | 0.04 | 0.00 | <0.001 | 0.08 |
| dPC2 | 0.00 | 0.00 | 0.09 | 0.07 | 0.00 | <0.001 | 0.03* |
| dPC3 | 0.00 | 0.00 | 0.04 | 0.05 | 0.00 | <0.001 | 0.42 |

<sup>a</sup> *Post hoc* Games-Howell analysis did not show any statistically significant differences between any of the force fields.

## Discussion

The use of robust statistical methods as outlined in the supporting materials^90-102^ have allowed us to systematically compare and document the effects of force field selection on the sampled conformational space of the inherently disordered protein Aβ(1-40) dimer in water at physiological temperature (310 K). Although, atomistic MD simulations have even been able to accurately predict the folded state of globular proteins, intermediate states are not consistent with experimental data and simulations have been unable to accurately reproduce melting curves and other temperature dependent behavior.^113-122^ The problems of inaccurate prediction of intermediate folded states and poor reproduction of melting curves arises from the overestimation of the steepness of the potential energy basins for protein-protein interactions.^34,109,129^ While force field depended protein-protein interactions tend to overestimate conformational stability, protein-water interactions tend to give rise to conformations that are overly compact compared to experimental data.^107,109,125^ These deficiencies pose significant problems when MD simulations are used for to study properties of inherently disordered peptides and proteins giving rise to unrealistic secondary structure sampling and stabilization.^34-39^

Determination of simulation equilibration and the length of time necessary to ensure adequate ensemble averaging are difficult and controversial. We have opted for a multifaceted approach, reviewing the Cα-trace configurational entropy; intra-monomer, inter-monomer, and dimer RMSIP; <P_2_> as a function of time; <R_g_> as a function of time; DSSP secondary structure as a function of time; and sampled ϕ,ψ dihedral angle space at 50 ns intervals (Figures 1-3 and Figures S1- S3 in the SI), as being a reasonable method of determination.^26^ Based on our results, all the systems clearly reached equilibration within 100 ns and the conformational sampling should occur from 100 to 300 ns.

The configurational entropy plateau values for the low temperatures (300 to 314 K) were similar for the AMBER-ILDN, AMBER-NMR, and the previously published CHARMM36 trajectories.^26^ There was significant spread in the plateau values for AMBER-ILDN* and to a lesser extent CHARMM22* however with significant difference in the rate of rise to the plateau values taking almost the entire 100 ns for AMBER-ILDN* compared to <25-50ns for CHARMM22*. The previously noted entropy inversion that occurred for the 310 K trajectory with decreased configurational entropy compared to other simulated temperatures for CHARMM36 was not noted for the other force fields.^26^

The results for <P_2_>, <R_g_>, and CCS are in general agreement with previously published results for Aβ(1-40) and other inherently disordered peptide systems. The values of <P_2_> are consistent with a fluctuating nematic droplet with a high fraction of β-sheet secondary structure.^65,66,103^ Despite only minor difference in the calculated values (Table 1) all are statistically significantly different as demonstrated by Welch’s ANOVA and *post-hoc* Games-Howell test. The calculated values of <R_g_> and CSS are consistent with ensembles of conformations that are slightly larger than experimental results (Table 1 and Table S1). These results consistent with the expected results for simulations of inherently disordered peptides and are both force field and water model dependent.^107,109,124^

Results obtained with AMBER-ILDN, AMBER-ILDN*, AMBER-NMR, and CHARMM22* demonstrate an increased probability of sampled αR-helical conformational space (Figures 2, 3 and 7; and Table 1). Although none of the compared force fields and simulations were able to reach statistical significant accuracy when compared to CD (Table S2), there are some significant similarities (Table 2). The fraction of predicted α-helix was similar between AMBER-ILDN* and CHARMM22* and the fraction of predicted turn/coil/bend was similar between AMBER-ILDN and CHARMM22*. The propensity for force fields of the AMBER family to over-stabilize and predict α-helical structures is well known.^123^ Our results for CHARMM22* differ significantly from those of Tarus and colleagues utilizing the same force field.^23^ While we used the DSSP method, Tarus and colleagues used the STRIDE algorithm. Although DSSP and STRIDE demonstrate excellent agreement for ordered peptide/protein structures, there may be significant differences when they are utilized for the analysis of inherently disordered peptides.^112^ Due to the difficulty in accurately assigning a compensatory factor to accommodate these differences, we assumed that the two results were approximately equal for our analysis. Our previously published results with CHARMM36 force field^26^ show a higher propensity to sample the β-sheet region of conformational space, which is still outside of statically significant similarity when compared to CD, but it confirms the AFM dynamic force spectroscopy results that both Aβ(1-40) and its residues 13–23 fragment form stable dimers in most likely antiparallel manner.^125,126^.

Examination of Cα-backbone stability and secondary structure (Figure 4) does demonstrate some degree of similarity between force fields. Areas of increased flexibility, other than that which is expected in the N- and C- terminal residues, tends to occur from residues 8-16 and 22-28 which roughly corresponds to regions of increased turn/bend/coil. The exception to this is the result for AMBER-NMR, which has a significant amount of α-helical secondary structure for residues 2-37. Those areas of increased stability correspond to the CHC and C-terminal hydrophobic cores. The associated secondary structure is, however, not consistently the predicted β-sheet-loop-β-sheet^20,21,27^ with AMBER-ILDN and AMBER-ILDN* performing better than CHARMM22*. Much of this disparity appears to be secondary to over sampling of α-helical content through the CHC, hydrophilic loop and C-terminal hydrophobic residues.

Although some of these differences in secondary structure are due to force field dependent protein-protein interactions, examination of SASA (Table 1 and Figure 5) per residue ∆E_Binding_ (Figure 8) and the probability of finding a water molecule within 0.5 nm of the acidic and basic residues (Table 4) would suggest that the role of protein-water interactions cannot be ignored. Both AMBER-ILDN and AMBER-ILDN* demonstrate increased rSASA in the loop region while AMBER-NMR has no such peak. CHARMM22* demonstrates an opposite effect with decreased rSASA for the CHC, C-terminal hydrophobic cores and the loop region. This is consistent with the lowest amount SASA_Hydrophilic_ for any of the force fields. The per residue differences in ∆E_Binding_ are, however, very minor between individual force fields (Figure 8) with strong favorable contributions from the basic residues (Arg and Lys) and strong unfavorable contributions from the acidic residues (Asp and Glu). The differences become clear with examining the probability of finding water molecules within 0.5 nm of the acidic and basic residues. CHARMM22* is slightly less likely, by approximately 1%, to have a water molecule within 0.5 nm of an acidic residue and slightly more likely, by approximately 1%, to have a water molecule within 0.5 nm of a basic residue. This contrasts dramatically with our previously published results for CHARMM36^26^ that indicated that this force field is more likely, by approximately 5%, to have a water adjacent to a an acidic residue but relatively similar to CHARMM22* with regards to basic residues. The increased hydration of CHARMM36 is in keeping with previously published results for this force field.^107^

The effect of intermolecular protein-protein and protein-water interactions undoubtedly play a role in dimer formation. The contact maps shown in Figure 6 are much sparser than those previously published for CHARMM36.^26^ AMBER-ILDN, AMBER-ILDN* and CHARMM22* all reproduce the anti-parallel contacts from residues 17-21 and 30-36, however, the probability of contact is much lower and the resulting quaternary structures are much less stable. Much of this may be explained by the calculated values of ∆E_Binding_ for all four force fields (Table 3) which are unfavorable, with the highest value being CHARMM22*. Although the ∆E_Binding_ for AMBER-NMR is less than those of AMBER-ILDN and CHARMM22*, the extended α-helical conformation seen for this force field makes the hydrophobic dimer interaction much less likely to occur. This contrasts with the previously published ∆E_Binding_ for CHARMM36 of −93.56 kJ•mol^−1^ which indicates favorable intermolecular protein-protein interactions. This favorable value of ∆E_Binding_ is also consistent with the DCCM analysis (Figure 9) which demonstrates strong correlated motions between the two monomer chains for the CHARMM36 force field only while strong anti-correlated motions occur for the AMBER-ILDN, AMBER-ILDN*, and AMBER-NMR force fields and a minimal week positive correlation for CHARMM22*.

Despite differences in calculated properties, Cα-backbone stability, sampled secondary structure, intra- and intermolecular contacts, and resulting lowest energy quaternary structures (Figure 10) the degree of similarity in essential subspace sampling (Figure 11 and Figure 12) is remarkable. The nRMSIP analysis suggests that the greatest degree of difference is 10% with most differences between force fields being on the order of 4-5% with the largest degree of difference occurring between the CHARMM36 and AMBER family of force fields. The ANOVA for dPCA1-3 would suggest however that these differences are not significant (Table 5).

## Conclusions

It is clear that the currently tested force fields (AMBER-ILDN, AMBER-ILDN*, AMBER-NMR and CHARMM22*) do not perform optimally for the simulation of an inherently disordered peptide dimer system. It is also clear from our own work with CHARMM36 as well as that of Raucher *et al.* that continued refinement of force field parameters is necessary.^26,34^ The MacKerell lab recently published a refinement of the CHARMM36 force field and associated water molecule (CHARMM36m and TIP3Pm) optimized for the simulation of inherently disordered peptides.^108,127^ The associated force field has modifications of the CMAP potential to correct backbone dihedrals and increasing the depth of the water hydrogen atom’s Leonard-Jones potential making the dispersion part of the protein-water interactions more favorable. There have also been significant modifications of the AMBER family of force fields with publication of AMBER-FB15 and associated TIP3P-FB water molecule which has also been optimized to simulate inherently disordered peptides as well as to accurately reproduce temperature dependent conformational behavior.^128^ AMBER-FB15 force field uses optimized dihedral angle potentials and 1-4 electrostatics to improve conformational sampling. Although both CHARMM36m and AMBER-FB15 demonstrate superior performance for simulation of inherently disordered peptide systems, it is unclear how they will perform for dimer or higher oligomer systems since the intermolecular peptide-peptide forces have not been optimized.^127,128^ Further work will need to be done on force field optimization order to obtain accurate results for these complex systems.

## Supporting Information Available

Supporting Material includes detailed expansion of the methods section describing the statistical methodology utilized and additional analyses of trajectories.

## Notes

The authors declare no competing financial interests.

## Acknowledgments

Authors thank to Dr. Yuguang Mu for providing his FORTRAN program for the dPCA analysis. The REMD simulations were completed using the Crane cluster at the Holland Computing Center of the University of Nebraska, Lincoln.

